# Structural and developmental principles of neuropil assembly in *C. elegans*

**DOI:** 10.1101/2020.03.15.992222

**Authors:** Mark W. Moyle, Kristopher M. Barnes, Manik Kuchroo, Alex Gonopolskiy, Leighton H. Duncan, Titas Sengupta, Lin Shao, Min Guo, Anthony Santella, Ryan Christensen, Abhishek Kumar, Yicong Wu, Kevin R. Moon, Guy Wolf, Smita Krishnaswamy, Zhirong Bao, Hari Shroff, William Mohler, Daniel A. Colón-Ramos

**Author notes:** Correspondence to: Daniel A. Colón-Ramos, Ph.D., Department of Neuroscience, Department of Cell Biology, Yale University School of Medicine, 333 Cedar Street, SHM B 163D, New Haven, CT 06510.

## Abstract

Neuropil is a fundamental form of tissue organization within brains^1^. In neuropils, densely packed neurons synaptically interconnect into precise circuit architecture^2,3^, yet the structural and developmental principles governing nanoscale precision in bundled neuropil assembly remain largely unknown^4–6^. Here we use diffusion condensation, a coarse-graining clustering algorithm^7^, to identify nested circuit structures within the *C. elegans* cerebral neuropil (called the nerve ring). We determine that the nerve ring neuropil is organized into four tightly bundled strata composed of related behavioral circuits. We demonstrate that the stratified architecture of the neuropil is a geometrical representation of the functional segregation of sensory information and motor outputs, with specific sensory organs and muscle quadrants mapping onto particular neuropil strata. We identify groups of neurons with unique morphologies that integrate information across strata and that create a sophisticated honeycomb-shaped scaffold that encases the strata within the nerve ring. We resolve the developmental sequence leading to stratified neuropil organization through the integration of lineaging and cell tracking algorithms with high resolution light-sheet microscopy, and reveal principles of cell position, migration and hierarchical outgrowth that guide neuropil organization. Our results uncover conserved design principles underlying nerve ring neuropil architecture and function, and a pioneer neuron-based, temporal progression of outgrowth that guides the hierarchical development of the layered neuropil. Our findings provide a blueprint for using structural and developmental approaches to systematically understand neuropil organization within brains.

---

To elucidate the structural and developmental principles that govern neuropil assembly, we examined the *C. elegans* nerve ring, a cerebral neuropil composed of 181 neurites^3^. The lineage, morphology and synaptic connectivity of all 181 neurites is known ^3,8^. While connectomic analyses have revealed network principles and circuit motifs^9–22^, we still lack an understanding of the design principles that underlie nerve ring neuropil architecture and function, and the developmental sequence that forms this functional architecture.

## Computational detection of a hierarchy of neurite bundling in the neuropil

To systematically dissect the organization of the nerve ring neuropil we quantified over 100,000 instances of neurite-neurite contact^22^ for each of two published *C. elegans* electron microscopy connectome datasets^3^, thereby generating two replicate contact profiles for all neurites within the nerve ring (**Fig. 1a**). Neurite placement is a major determinant of synaptic connectivity^23^. We therefore selected contact profiles instead of synaptic connections to reveal relationships among all neurons, both functional and structural.

**Figure 1:**
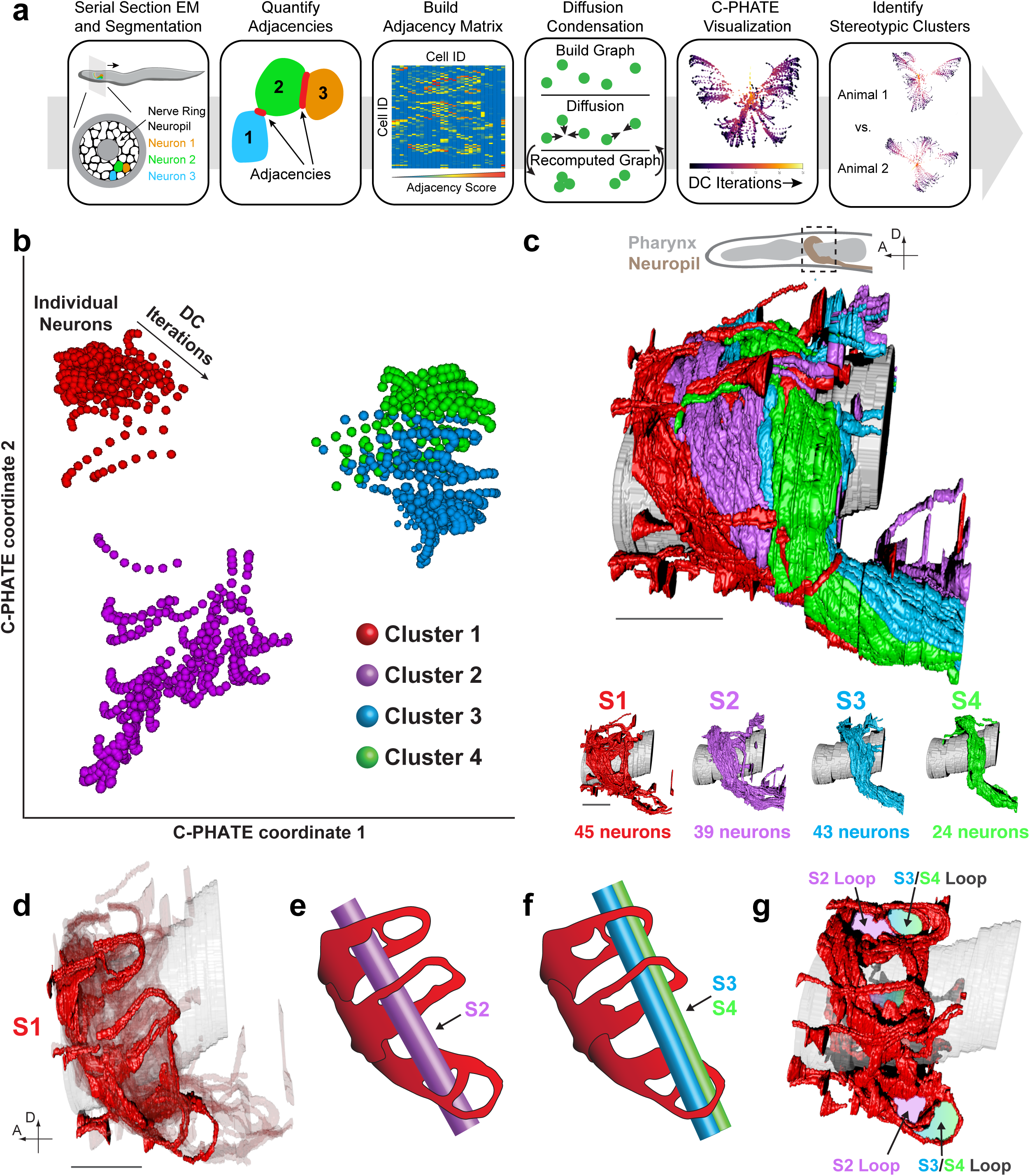
Computational detection of hierarchical neurite bundling in the *C. elegans* neuropil. **a,** Pipeline for analyses of the *C. elegans* neuropil. Serial section EM data^3^ was used to define all neuron-neuron contacts^22^. An adjacency matrix of neuron-neuron contacts was generated and analyzed via Diffusion Condensation (DC)^7^, and the output visualized via C-PHATE^30^. Output for two animals was quantitatively compared and stereotypic clusters and outliers identified. **b,** C-PHATE plot of DC analysis for a larval stage 4 (L4) animal (DC iterations 1-11 shown). Individual neurons are located at edges of the graph and condense as they move centrally. The four super-clusters identified are individually colored. (Supplementary Videos 1-2) **c,** Volumetric reconstruction of the L4 *C. elegans* neuropil (from EM serial sections^3^) with the 4 strata individually colored. Below are representations of individual stratum. Scale bar is 5 µm; Extended Data Fig. 1c-h, Supplemental Video 3. **d,** Volumetric reconstruction of scaffolding neurons (highlighted) in S1. Scale bar is 5 µm. **e,** Schematic representation of (**d**) with the trajectory of the S2 bundle (in **c**) through the specific loops formed by the scaffold. **f**, Like (**e**), but with the S3/4 bundle. **g,** The honeycomb-like scaffold formed by 32 of the 45 neurons in S1, with loops colored for the positions of the encased strata (in **c**). Extended Data Fig. 3 and Supplemental Videos 4-5.

To the adjacency matrix of contact profiles we applied “Diffusion Condensation” (DC^7^), a coarse-graining algorithm that iteratively groups neurons based on the quantitative similarity of each neuron’s profile of adjacencies (**Fig. 1a**). Unlike many other clustering algorithms^24–29^, DC condenses data without assumptions regarding underlying data structure, and by iteratively moving datapoints closer to their diffusion neighbors. This dynamic process unveils data relationships at varying scales of granularity, from cell-cell interactions to circuit-circuit bundling.

To visualize DC cluster relationships we developed C-PHATE, an extension of PHATE^30^ capable of generating a 3D representation of the multigranular structures in the DC hierarchical tree (**Fig. 1a,b**; see Methods). Quantitative comparisons of DC/C-PHATE results revealed similar clustering behaviors between the two separately analyzed nerve-ring reconstructions (Adjusted Rand Index (ARI) of 0.7; **Extended Data Fig. 1a,b**), consistent with the long-accepted qualitative descriptions of the stereotyped structure for the *C. elegans* nerve ring^3,31^, and recent analyses of neurite adjacencies^22^. Examination of the cluster patterns through the iterations of DC revealed known cell-cell interactions and behavioral circuits (**Fig. 1b,c; Extended Data Fig. 2a,b**;^32–39^), consistent with behavioral and connectomic studies^10,19,20^. The multigranular outputs of the DC/C-PHATE results enabled understanding of the cell-cell interactions within the context of functional circuits, and the functional circuits within the context of the larger bundles of the neuropil.

Comparisons of the modularity score, a measure of cluster robustness^40,41^, was highest when the majority of neurites were grouped into four super-clusters (**Fig. 1b; Extended Data Fig. 1a,b; Supplemental Video 1,2**; see Methods). Overlaying our analysis of these four super-clusters (see Methods) onto the anatomy of the nerve ring revealed that they correspond to four distinct and tightly grouped bundles within the greater neuropil, circling the pharynx isthmus and stacked along the anterior-posterior axis of the animal. These clusters are herein called S1, S2, S3 and S4 for Stratum 1 etc. (**Fig. 1c; Extended Data Fig. 1c-h; Supplemental Video 3** show details of these structures and single-cell identities). The stratified organization of the nerve ring neuropil, resolved here at single neurite level, is reminiscent of laminar organizations in the *Drosophila* nervous system^42,43^, and in the vertebrate retina and the brain cortex^44,45^.

We observed that not all computationally clustered neurites followed simple bundled paths through the neuropil. In Strata 1 (S1), 32 anterior sensory neurons project perpendicular to the nerve ring bundle before making a 180° curl and returning to the anterior limits of the neuropil, where they terminate as synaptic endplates (**Fig. 1d; Extended Data Fig. 3a-d;**^3,46^). These uniquely shaped S1 neurons looped at the borders of the S2/S3/S4 bundles. The anterior loops encase ∼90% of S2, and the posterior loops encase ∼84% of S3 and 100% of S4 (**Fig. 1d-g; Extended Data Fig. 3e-k; Supplemental Video 5; Supplemental Table 1**). We found that the 32 axons form a six-fold symmetrical honeycomb structure along the neuropil’s arc, creating a scaffold that encapsulates the different strata (**Fig. 1g; Extended Data Fig. 3e-h; Supplemental Video 4**).

## The architecture of the neuropil reflects functional segregation of distinct sensory information streams and motor outputs

*C. elegans* neurons can be divided into three categories based on structural and functional properties: sensory neurons, interneurons, and motor neurons^19,47^. Sensory neurons represent the first layer of information sensing and encoding in the nervous system. Therefore, to understand the functional anatomy of the nerve ring, we examined axonal positions of the head sensory neurons within the anatomy of the nerve ring neuropil.

There are two main sensory classes of neurons at the anterior buccal tip of the animal: papillary and amphidial sensilla^46^. While these two neuron classes are in close proximity, they are distinguished by distinct dendritic sensory endings, which are proposed to represent distinct sensory modalities^46,48^. Both classes of neurons project axons into the neuropil to transduce sensory information onto the nerve ring^46,48^. We observed that the papillary sensilla neurons exclusively project axons to the anterior bundle S1 (**Fig. 2a-c**), while the amphidial axons project to the S3 or S4 bundles (**Fig. 2a,b,d**). No papillary or amphidial axons project onto the S2 bundle. Our findings reveal that sensory neuron classes, historically defined by the morphology of dendrites^46^, map onto specific strata of the *C. elegans* nerve ring, suggesting a functional segregation of sensory information and processing in the neuropil’s layered structure.

**Figure 2:**
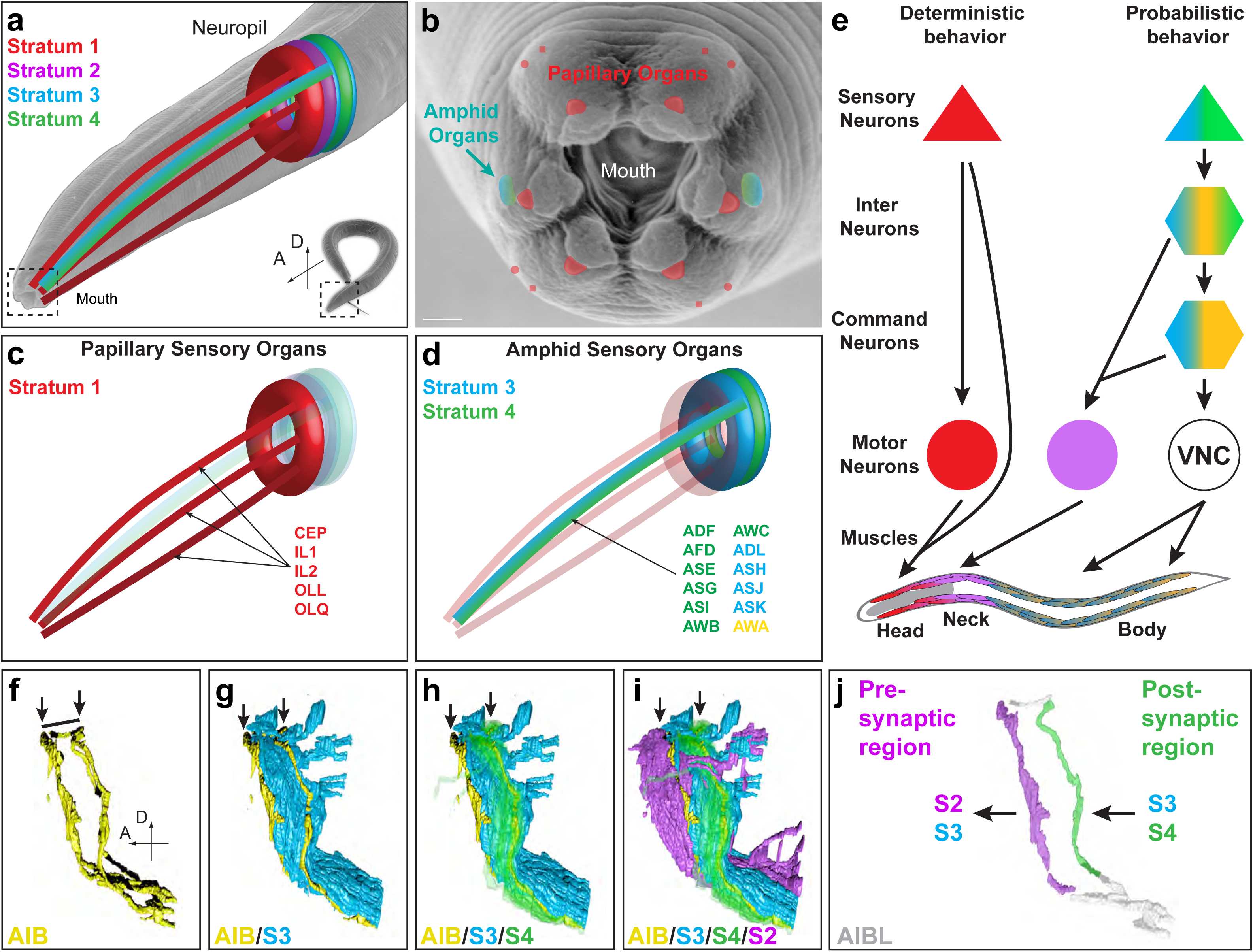
Neuropil architecture reflects functional segregation of sensory and motor outputs. **a,** Representation of head sensory organs in *C. elegans* in the context of the four strata, colored as in (Fig. 1). Representation projected over a scanning EM of *C. elegans* (inset lower right), enlarged to show position of sensory organs and strata in the head (Image from WormAtlas, produced by and used with permission of Ralf Sommer). **b,** Representation of head sensory organs, projected over a scanning EM of the *C. elegans* mouth (correspond to dashed box in lower-right of (**a**); image produced by and used with permission of David Hall. Mouth sensilla colored according to their strata assignment. Scale bar is 1 µm**. c,** Schematic of papillary sensory organs’ trajectories from mouth to neuropil. All papillary neurons (six-fold symmetry organs in (**b**)) cluster into S1 bundle (red). Individual neuron classes listed in lower-right. **d,** As (**c**), but with amphidial sensory organs. AWA is yellow because it belongs to ‘unassigned’ group (see Extended Data Fig. 4 e,f,q). **e,** Model of functional segregation of distinct sensory information streams and motor outputs within the neuropil. Papillary sensory information is processed in shallow circuits in the S1 bundle and innervate head muscles. Amphid sensory information is segregated to S3/S4, where interneurons process and link to body muscles (via command interneurons) and neck muscles (via motor neurons in S2). Shallow papillary circuits are associated with deterministic behaviors, while amphid sensory responses are associated with probabilistic/plastic behaviors^35,49,50,54^. Circuit units colored according to strata (S1-Red, S2-Purple, S3-Blue, S4-Green, Unassigned-Yellow); Extended Data Fig. 2c. **f-i,** Volumetric reconstructions of the unassigned (yellow), ‘rich-club’ AIB interneurons^18,21^ in the context of nerve ring strata. Arrows indicate the two segments of AIB that border strata. AIB proximal neurite borders S3/S4 (**g-h)**, shifts along the A-P axis, and the distal neurite borders S2/S3 (**h-i**). Line in (**f**) indicates AIB shift along the A-P axis to change strata. In (**h**) S4 is transparent to show AIB bordering of S3 and S4; Extended Data Fig. 2d,e and Supplemental Video 6. **j,** Volumetric rendering of AIBL. AIB is a unipolar neuron that displays polarity in the position of its pre and postsynaptic sites along the neurite. The synaptic position and the morphology of AIB allows it to receive sensory information from the S3/S4 amphid strata and transmit it to the S2, motor-neuron rich strata (see Extended Data Fig. 2d,e).

Detailed examination of the distribution of sensory neurons, interneurons and motor neurons further revealed design principles of neuropil functional anatomy. Within S1, the papillary sensory cells, which are mechanosensory or polymodal, control head withdrawal reflex behaviors^49,50^. This reflex behavior, from sensory neuron to motor output, maps onto the S1 bundle. Consistent with the deterministic nature of the behavior, most neurons in the S1 bundle are part of shallow circuits formed by papillary sensory cells connecting directly onto motor neurons or muscles^3,50^. Indeed, IL1 and URA sensory neurons form direct sensory-to-muscle synapses within the S1 bundle^3^. Furthermore, neuromuscular synapses from S1 form exclusively onto head muscles^3,46,48^ (**Fig. 2e; Extended Data Fig. 2c**).

Interestingly the S1 circuits retain the four-fold symmetry of the papillary sense organs at the interneuron, motor neuron and head neuromuscular synapse level^3,46,48^. Topographic maps—the ordered projection of sensory information onto effector systems like muscles—is a fundamental organizing principle of brain wiring across sensory modalities and organisms^51–53^. We find that the S1 circuits in *C. elegans* display a topographic map organization, from the primary sensory layer to the motor output representations.

In contrast we found that the amphid sensory axons, which are associated with plastic behaviors^35,54^, are located in the S3 and S4 layers. The S3 and S4 layers also contain interneurons, but lack motor neurons. The primary and secondary interneurons in the S3 and S4 layers connect with motor neurons in the S1 and S2 layers (to innervate head and neck muscles) or to command neurons (that connect to motor neurons which innervate body-wall muscles) (**Fig. 2e; Extended Data Fig. 2c**). Therefore, information streams emerging from the amphid sensory axons segregate onto S1 and S2 (to control head/neck muscles) or onto command neurons in S3 (to control body wall muscles). Our findings are consistent with cell ablation and behavioral studies that identified modular navigation strategies in response to amphid sensory signals^55,56^ and with connectomic studies that demonstrated the modular nature of these motor outputs ^19,22^. Deterministic reflex behaviors (like the head withdrawal reflex) or probabilistic plastic behaviors (like the chemotaxis behaviors) differentially activate distinct motor strategies in response to sensory information^55^. Our observations uncover the topological representations of these behavioral strategies in the architecture of the neuropil, and reveal fundamental design principles in nerve ring layered structure, from sensation to motor outputs.

## Unassigned cells correspond to ‘rich-club’ interneurons that connect across strata

The four major neurite strata account for 151 of the 181 total neurons in the nerve ring (83%). To further understand the nerve ring’s structure, we carefully examined the remaining 30 neurons that were not consistently assigned to a stratum (see Methods, herein called “unassigned”). These neurons fell into three broad categories: a) Neurons with simple, unbranched processes between boundaries of two strata (6 neurons); b) Neurons with unique morphologies that cross strata, including neurite branches projecting into multiple strata or single neurites that project perpendicularly across strata (21 neurons); c) Neurons with sparse segmentations (3 neurons) (**Extended Data Fig. 4**). Comparisons of our findings with previous connectomic studies revealed that six of the neurons identified by our analyses as “unassigned” belong to a group of 14 highly interconnected interneurons previously defined as “the rich-club”^18,21^. “Rich-club” refers to a conserved organizational network property seen for information-processing systems (such as brains) in which highly interconnected hubs (neurons) link segregated modules^57^. To uncover the roots of rich-club organization in the context of the architecture of the neuropil, we examined the cell biology of these unassigned neurons in the context of the strata.

We focused on unassigned, ‘rich-club’ interneuron AIB (detailed analyses of remaining unassigned and ‘rich-club’ neurons in **Extended Data Fig. 4**). We observed that the cell morphology, cell polarity and cell position of the AIB interneurons are precisely positioned to receive inputs from S3/4, and to transduce outputs onto S2/3, linking these modular strata. AIB neurite is positioned at the border between S3 and S4, with a perpendicular shift of precisely the width of S3, and a distal neurite at the border of S2 and S3 (**Fig. 2f-i; Supplemental Video 6**). AIB synaptic positions are similarly designed to connect the amphid sensory-rich S3/4 bundles with the motor neuron-rich S2 bundle (**Fig. 2j; Extended Data Fig. 2d,e**). This architecture is consistent with AIB’s known importance in processing amphid-derived sensory stimuli to mediate locomotory strategies^55,58–60^. We note that ‘rich-club’ interneuron AVE has a similar morphology to AIB, but with its neurite anteriorly displaced one stratum (**Extended Data Fig. 4a-d; Supplemental Video 7**). We observed that other “rich-club neurons”, such as RIB and RIA, as well as other “unassigned” neurons, such as AIZ, similarly shift across strata to switch neighborhoods, revealing design principles of inter-strata functional connectivity (**Extended Data Fig. 2f and 4a-d,h-u**). Notably, the nematode *P. pacificus*, which shared a common ancestor with *C. elegans* over a 100 million years ago, has a similar neuropil architecture, including the presence of rich-club interneurons which cross strata (like AIB) and the honeycomb scaffolding structure from S1^61^. The evolutionary conservation of these design principles in neuropil architecture suggests that these conserved structural motifs might underpin important aspects of nerve ring function.

Overall, our analyses reveal design principles at varying degrees of granularity— from single ‘rich-club neuron’ morphologies that functionally bridge different strata, to layered bundles that segregate sensory-motor information onto topological maps. These design principles are important organizational units in neuroscience—‘rich-clubs’ in the context of brain networks^18,21^, laminar organization in the context of brain structures^42–45^ and topological maps in the context of vertebrate sensory systems^51–53^. We map them in the context of the neuropil architecture and resolve them for *C. elegans* at cellular and synaptic scales.

## Layered neuropil architecture is influenced by early segregation of neuronal somas via cell migration

Knowledge of neuropil structure provides a unique opportunity to understand the developmental logic leading to its assembly. To visualize the developmental sequence leading to nerve ring assembly, we established an integrated protocol that enables long-term, four-dimensional imaging of living *C. elegans* embryos, with isotropic spatial resolution^62^. Our system yields minute scale high-resolution 4D data sets, enabling simultaneous tracking of nuclei and neuronal structures^63–68^, which can be analyzed to correlate cell-lineage identities with developmental dynamics.

The lineage identities for *C. elegans* are known and invariant^8^, enabling us to rigorously establish the relationship between individual neuron lineage identity, its nuclear positions during embryonic development and neurite positions in the nerve ring strata. Time lapse datasets of lineaged and tracked positions for all 580 cells in a developing embryo (from ∼70 to 430 minutes post fertilization (mpf)), were systematically examined for birth order, neuronal precursor positions and lineage identity in the context of the future strata. Although it has been previously hypothesized that lineage-dependent neuronal soma positions might influence the outgrowth of neurites into neighborhoods^69^, we could not detect any relationships between lineage, soma position upon the last cell division and the neurite position within the neuropil strata (**Fig. 3a; Extended Data Fig. 5**).

**Figure 3.**
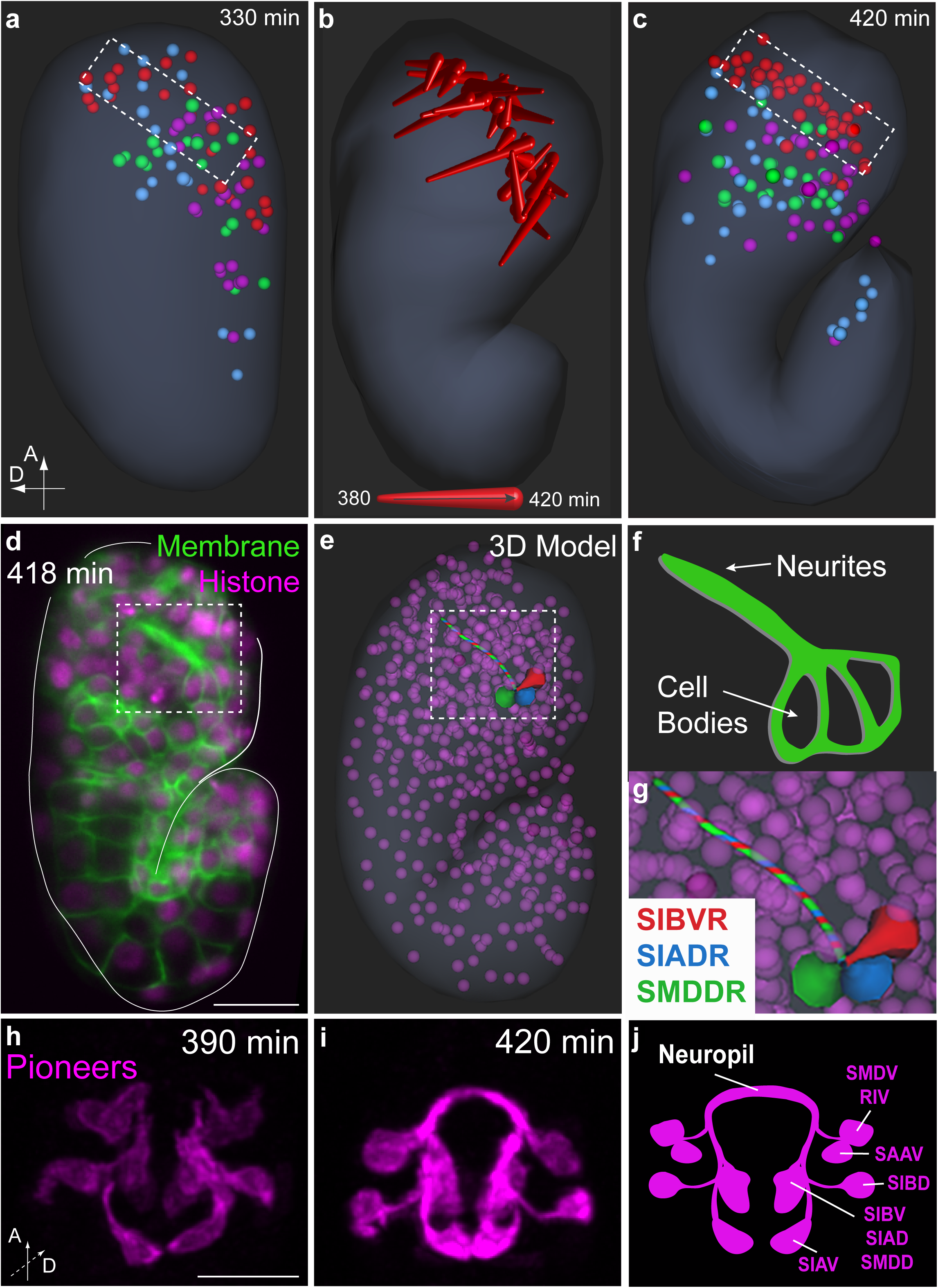
Segregation of S1 neuronal somas via cell migration, and pioneering S2 outgrowth events, influence layered neuropil assembly. **a,** Representative model of all neuronal soma positions within the embryo (generated using the WormGUIDES atlas^66^) at 330 min post fertilization (mpf). Somas are colored to show their eventual neurite strata assignment. **b,** 3D depth trajectories of S1 cells movement between 380-420 mpf. **c,** As, (**a**), but at 420 mpf. Dashed box in (**c**) as in (**a**), represents the final position of the migrating S1 neurons in the anterior of the embryo head; Extended Data Fig. 6a-h; Supplemental Video 8. **d,** Comma stage embryo (418 mpf) labeled with GFP membrane marker (promoter nhr-2) expressed in all cells and mCherry Histone (to identify nerve ring outgrowth events and cell positions/lineage, respectively). Image is single z-slice from one diSPIM arm. Scale bar is 10 µm; Extended Data Fig. 6i-n, and Supplemental Video 9. **e,** 3D model of the three cells observed in (**d**) (made with the WormGUIDES atlas^66^). **f,** Cartoon of the inset of the 3 cells observed in (**d**). **g,** Enlargement of inset (**e**), with early outgrowing cells identified via lineaging, listed and colored to highlight cellular locations. **h-j,** Time-lapse of the outgrowth dynamics of pioneer neurons (labeled with Plim-4::membrane-tethered::GFP) and schematic. Images are deconvolved diSPIM maximum intensity projections; Extended Data Fig. 6o-z, and Supplemental Video 10. Scale bar (10 µm) applies to panels (**h,i**). Timing is mpf for all images.

Post-mitotic cells in *C. elegans* embryos are capable of short migrations during organ morphogenesis^8,70^. To understand the role of global cell migrations in the emerging nerve ring, we generated a developmental atlas (called WormGUIDES) that represents the 3D positions of all cells in the developing embryo through time^66^. Quantification of the positions of individual neurons (belonging to specific strata) in the context of the spatio-temporal dynamics of embryo morphogenesis revealed coordinated cell movements that segregated, and co-located, cell bodies of future strata (**Fig. 3a-c**). For example, cell bodies of neurons which later project onto the S1 bundle migrated and co-located to the anterior part of the embryo head, while cell bodies of neurons which later project onto the S2-S4 strata migrated to segregate at the posterior part of the head (400-430 mpfs, prior to neurite outgrowth; **Fig. 3a-c; Extended Data Fig. 6a-h; Supplemental Video 8**). The positions adopted during embryogenesis persist until adulthood, and for the future S1 strata, relate to the cellular morphologies of posteriorly-projecting axonal structures within the anterior bundle of the nerve ring^3,46,48^.

In vertebrate embryogenesis, migration of waves of neurons help organize the layered architecture of the retina and brain cortex^71,72^. Our systematic examination of neuronal lineage, birth timing and global migrations reveal that while birth positions do not directly correlate with the stratified architecture of the neuropil, co-location of neuronal cell bodies through global migration serve as an initial organizing principle that corresponds to the antero-posterior layering of the neuropil, and the functional segregation of the sensory-motor architecture in the nerve ring.

## A temporal progression of outgrowth, guided by pioneer neurons, results in the hierarchical development of the layered neuropil

To elucidate additional organizing principles that contribute to neuropil assembly, we imaged the outgrowth dynamics of neurites during formation of the neuropil in embryos by expressing a ubiquitous membrane tethered GFP (**Extended Data Fig. 6i-n; Supplemental Video 9**). At approximately 400 minutes post fertilization (mpf) we noticed 6 cells sending projections into the area of the future nerve ring (**Fig. 3d,f**). Using simultaneous mCherry::histone imaging and the cell lineage-tracking tools StarryNite^63^ and AceTree^73,74^, we identified those 6 cells as 3 bilateral pairs of neurons: SIAD, SIBV, and SMDD (**Fig. 3e,g**). To confirm their identities, we co-labeled embryos with ubiquitous membrane-tethered GFP and a cytoplasmic Plim-4::mCherry reporter gene (previously lineaged^66^ to express in SIAD, SIBV, and SMDD (**Extended Data Fig. 6o-r**) and an additional 5 pairs of neurons (RIV, SAAV, SIAV, SIBD, SMDV)). We observed that all 16 labeled cells extend bright red neurites into the future neuropil region at 400 mpf (**Fig. 3h-j; Supplemental Video 10**).

The identified neurites belong to a nerve bundle located in the central strata, S2 (with the exception of neuron SIBV, which, while part of the same bundle, is considered by our analyses as an “unassigned neuron” due to its morphology, see **Extended Data Fig. 4e,f,o**). When compared to a pan-neuronal marker for nerve ring development, these cells displayed the earliest outgrowth events observed for the neuropil (**Extended Data Fig. 6s-z; Supplemental Video 10,11**). Given the central position of these neurons in the neuropil and their early outgrowth during development, we hypothesized that they might be pioneers for the developing neuropil. Consistent with our hypothesis, a previous genetic study examining nerve ring placement proposed that two of the neurons also identified in our study, SIA and SIB, position the nerve ring neuropil during development^75^. Additionally, a study examining the role of glia in nerve ring development demonstrated that ablation of SIA and SIBV resulted in outgrowth defects of interneuron AIY and the amphid commissure, suggesting they were important for development of neurons into the nerve ring^76^.

To examine this hypothesis, we adapted an *in vivo* split-caspase ablation system^77^ that killed these 16 neurons during embryonic neurodevelopment (**Extended Data Fig. 7a-f**). We then systematically monitored neuropil formation using a pan-neuronal reporter (**Supplemental Video 12**). Ablation of these putative 16 pioneering neurons resulted in larval stage 1-arrested animals with severely disrupted neuropils, as determined by total embryonic neuropil volume and structure (mean control neuropil volume for wild type was 136.6 µm^3^ at 490 mpf, while mean ablation neuropil volume at 490 mpf was 43.6 µm^3^; **Fig. 4a-c; Extended Data Fig. 7g,h**). Systematic examinations of cell-specific promoters that label neurons in the four strata revealed that the outgrowth of all examined neurons was affected upon ablation of the purported pioneer neurons (**Fig. 4d-f; Extended Data Fig. 8**). Notably, ablation of the purported pioneers in the S2 bundle similarly affected the outgrowth phenotypes for neurites belonging to the S3, S4 bundles and “unassigned” neurons (ASH, AIY and AIB neurons respectively; **Extended Data Fig. 8c,h,v,y**). In all these cells, neurites paused indefinitely near the ablated pioneer somas. Additionally, we found that the S2 pioneers are centrally positioned within the neuropil along both the anteroposterior and dorsoventral axes (**Fig. 4g-i**). Our findings demonstrate that the nerve ring develops via a hierarchical process, beginning with a subset of centrally-located S2 pioneer neurons.

**Figure 4.**
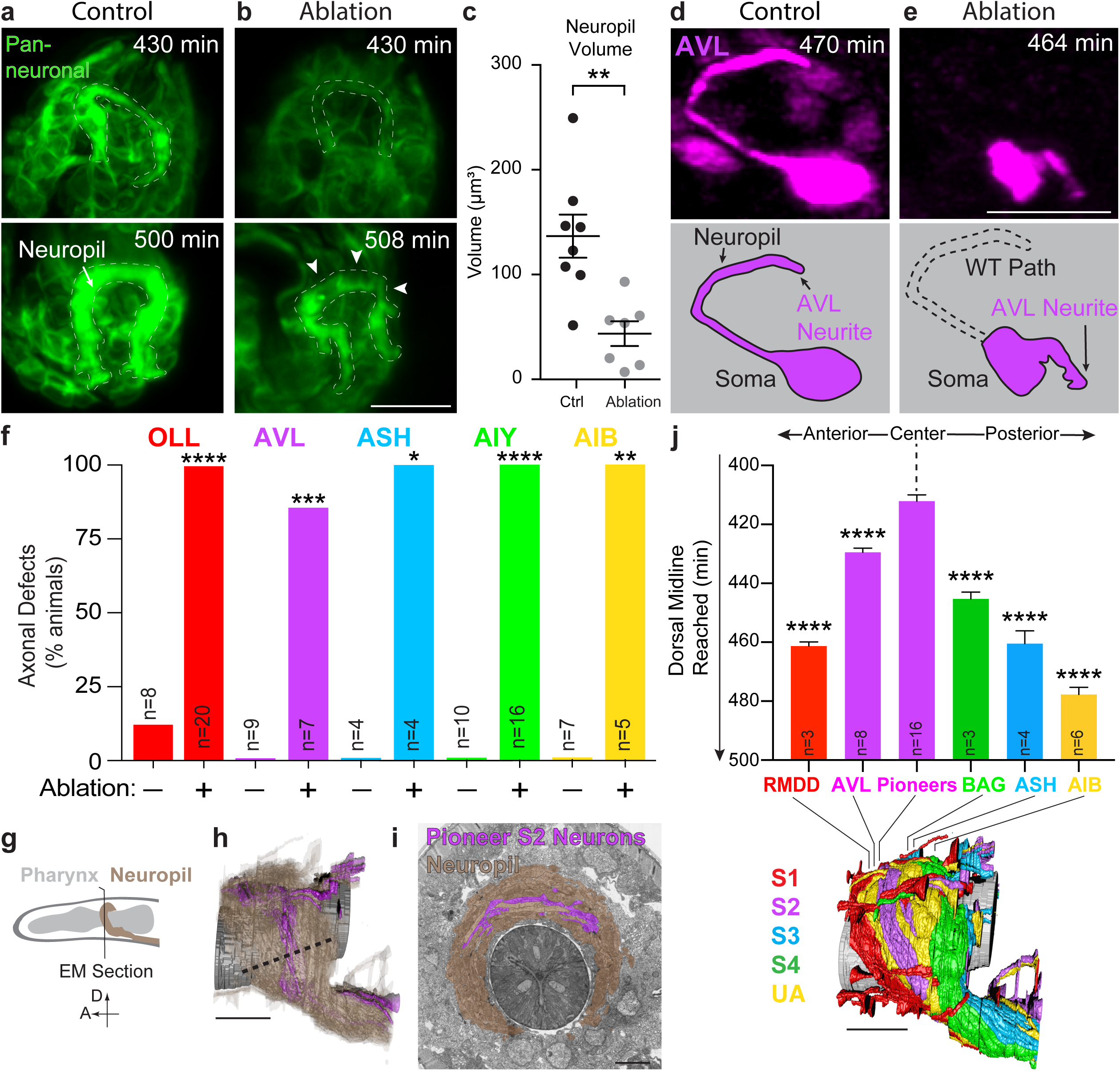
A temporal progression of outgrowth, guided by centrally located S2 pioneer neurons, results in the hierarchical inside-out development of the layered neuropil. **a-b,** Neuropil development for control (**a**) or pioneer ablated (**b**) embryos monitored via pan-neuronal marker (Pceh-48::membrane-tethered::GFP) and light sheet microscopy^64,65^. Dashed lines represent the position of the wild type neuropil. Maximum intensity projections of data from one diSPIM arm; Supplemental Video 12. Scale bar (10 µm) applies to all panels. **c,** Quantification of neuropil volume for control or pioneer ablated animals. **p<0.01 by unpaired Student’s t test between Ctrl and Ablation. Error bars are SEM. Extended Data Fig. 7g-h. **d-e,** AVL development in control (**d**) or pioneer ablated (**e**) embryos monitored using an AVL-specific promoter (Plim-6::GFP). Dashed line depicts normal AVL outgrowth. Deconvolved diSPIM maximum intensity projections are shown (for other neurons analyzed see Extended Data Fig. 8); images in (**d**) same dataset as Supplemental Video 13. Scale bar (10 µm) applies to all panels. **f,** Quantification of neurite outgrowth for indicated neurons from each stratum in control and pioneer ablated animals. Color indicates the strata to which each neuron is assigned. ‘n’ is the number of embryos (AVL, ASH, AIB) or L1 animals (OLL, AIY) scored. *p<0.05, **p<0.01, ***p<0.001, ****p<0.0001 by Fisher’s exact test between Ctrl and Ablation for each neuron; Extended Data Fig. 8a-z. **g,** Schematic of worm head and neuropil with position of EM cross-section shown in (**h,i**). **h,** Volumetric reconstruction of the *C. elegans* L4 neuropil. S2 pioneers in purple, neuropil in brown, pharynx in grey. Dashed line indicates width of neuropil. Note that the S2 pioneers are centrally-located in the neuropil. Scale bar is 5 µm. **i,** Segmented serial section electron micrograph from^3^, with centrally-located pioneer neurons in purple, neuropil in brown, pharynx in dark grey (section corresponds to JSH z-slice 54). Scale bar is 2.5 µm. **j,** Analysis of dorsal midline outgrowth for neurons from each stratum. Below is a volumetric neuropil reconstruction with the terminal location, along the dorsal midline, of the examined neurons and their relative positions to the centrally-located pioneers. ‘n’ is the number of embryos scored. ****p<0.0001 by one-way ANOVA with Tukey’s post hoc analysis between pioneers and the representative neurons from each stratum. With Tukey’s post hoc analysis all pairwise neuron comparisons are significant (*p<0.05) except for between RMDD, BAG, and ASH. Error bars are SEM. Scale bar is 5 µm; Extended Data Fig. 9a-i’. Timing for all panels is mpf.

Next, we documented the developmental sequence of outgrowth in the context of the central S2 pioneers and the known neuroanatomy of the nerve ring. We achieved this by monitoring synchronized recordings of neurite outgrowth from individual neurons in each stratum in developing embryos, and then comparing their temporal outgrowth dynamics with their terminal location in the nerve ring relative to the pioneering neurites. We observed an ordered sequence of outgrowth events in which the timing of embryonic entry into the neuropil (but not the timing of outgrowth from the cell body) tightly correlated with the radial proximity of the examined neurite to the pioneer neurites (**Fig. 4j; Extended Data Fig. 9a-i’; Supplemental Video 13-15**). Notably, neurons from S4 were observed initiating outgrowth at 400 mpf, a similar time as the pioneering neurons (**Fig. 3h; Extended Data Fig. 9j**). But instead of entering the nerve ring like the pioneers, S4 neurites paused for approximately 30 minutes near the somas of the pioneering SAAV neurons and then continued outgrowth into the nerve ring. (**Fig. 3h,i; Extended Data Fig. 9j-q**). We note that the pausing point for the examined neurons correspond to a similar location of defective outgrowth stalling upon pioneer neuron ablations (**Extended Data Fig. 8c,h,v,y**). Our findings indicate that while the initial outgrowth events for many neurons occur simultaneously, neurites extend to, and pause at, nerve ring entry sites. A temporal progression of tiered outgrowth events into the nerve ring, which initially depends on centrally located pioneer neurons, then results in the layered development of the nerve ring neuropil.

The temporal sequence of outgrowth events into the nerve ring continues through late embryogenesis into larval stage four (L4), and follows the design principles described earlier for the neuropil structure. For example, outgrowth of ‘rich-club’ interneuron AIB initiates at 420 mpf and arrives at the dorsal midline at 480 mpf, after the architecture of the nerve ring strata is established (**Fig. 4j; Extended Fig. 9e-I’**). Similarly, the S1 neurites which result in the six-fold symmetrical honeycomb structure that encases the neuropil strata develop after the strata have been assembled. All of these outgrowth events were similarly dependent on the pioneer neurons for their correct development. The embryonic sequence of neurodevelopment uncovered here, as well as the known outgrowth events which occur during larval stages^78^, indicate the existence of a hierarchical sequence, coordinated through time, which result in the assembly of the layered architecture of the neuropil.

Lamination is a conserved principle of organization within brains^45^. Segregation of functional circuits into layers underpins information processing in sensory systems and higher order brain structures^44^. In this study we resolve these conserved features of brain organization in the context of the nerve ring, linking fundamental design principles of neuropil organization with the developmental processes that underpin their assembly.

We achieve this by categorizing neurons based on their structural relationships in the neuropil. We note that there are other ways of categorizing neurons distinct from the ones presented in this study—by anatomical location of functional units^3^, neurotransmitter expression^79^, or connectivity^11,19^, among others. Our categorization, which is not exclusive to other neuronal categories, reveal useful principles in neuropil function and development. The modular organization of the neuropil strata revealed by our analyses is important for understanding information processing organization underpinning behavior, and to elucidate conserved developmental principles that instruct precise layered assembly in nervous systems. Our findings also provide a blueprint regarding synergistic integration of structural connectomic analyses and developmental approaches to systematically understand neuropil organization and development within brains.

## Acknowledgements

We thank Scott Emmons, Steve Cook, and Chris Brittin for sharing the segmented EM datasets and adjacency analysis code^22^, and for helpful comments on this work. We thank Oliver Hobert for sharing the Pceh-48 promoter and advice. We thank Zheng Zhou for sharing a 2xFYVE containing plasmid^80^. We thank David Hall for advice and help with electron microscopy images. We thank Ralf Sommer for use of an electron microscopy image. We thank members of the Colón-Ramos lab for insightful comments during manuscript preparation. We thank the Caenorhabditis Genetic Center (funded by NIH Office of Research Infrastructure Programs P40 OD010440) for *C. elegans* strains. We thank the Research Center for Minority Institutions program, the Marine Biological Laboratories (MBL), and the Instituto de Neurobiología de la Universidad de Puerto Rico for providing meeting and brainstorming platforms. H.S. and D.A.C-R. acknowledge the Whitman and Fellows program at MBL for providing funding and space for discussions valuable to this work. Research in the D.A.C-R., W.M., H.S. and Z. B. labs were supported by NIH grant No. R24-OD016474. M.W.M was supported by NIH by F32-NS098616. Research in H.S. lab was further supported by the intramural research program of the National Institute of Biomedical Imaging and Bioengineering (NIBIB). Research in Z.B. lab was further supported by an NIH center grant to MSKCC (P30CA008748). Research in the D.A.C.-R. lab was further supported by NIH R01NS076558, DP1NS111778 and by an HHMI Scholar Award.

## Extended Data Figure Legends

**Extended Data Figure 1. Computational analysis of the DC clustering identifies 4 strata within the neuropil.**

**a-b,** Quantification of network modularity^41^ for DC analysis of L4 (**a**) and adult (**b**) animal. Network modularity was determined by creating an affinity graph for each iteration and then calculating modularity of that graph. Note that the highest modularity score was for the iteration with four clusters in the L4 and 6 cluster in N2U. Comparisons revealed that for N2U, there were four large clusters similar to JSH, and two smaller ones (see Methods). These iterations were used for subsequent strata analyses (see Methods) **c,** Volumetric reconstruction of the L4 *C. elegans* neuropil (from EM serial sections^3^) with the 4 strata and unassigned neurons individually colored. Above, schematic of worm head with nerve ring neuropil (dashed box). Location of EM sections displayed in (**d-g**) shown below (arrows and corresponding section numbering). Scale bar is 5 µm. **d-g,** Segmented serial section electron micrographs from^3^, neurons colored as in (**c**). Original EM slices 41, 92, 152, 206 shown in (**d-g)** respectively. Scale bar (2.5 µm, in **d**) applies to panels (**d-g**). **h,** Neuronal classes in the 4 strata, and the ‘unassigned’ group. Number to the right of each neuron represents total neurons for each class, for all 181 neurons in the embryonic nerve ring neuropil.

**Extended Data Figure 2. Examination of behavioral circuits in the DC/C-PHATE analyses**

**a,** C-PHATE plot of DC analyses for a larval stage 4 (L4) animal, with known behavioral circuit locations highlighted. The four super-clusters identified are colored accordingly (C1-Red, C2-Purple, C3-Blue, C4-Green). **b,** Enlargement of inset from (**a**) displaying the condensation of a group of neurons corresponding to the mechanosensation circuit^37,56,81,82^. Neurons with their names in filled blue boxes are members of the anterior body mechanosensation circuit, neurons with their names in outlined blue boxes (BDUL and BDUR) guide the formation of the circuit during development^82^. **c,** Model of functional segregation of distinct sensory information streams and motor outputs within the neuropil in the context of known behavioral circuits. Papillary sensory information is processed in shallow circuits in the S1 bundle and innervates head muscles. Amphid sensory information is segregated to S3/S4, relayed to interneurons and then to body wall muscles (via command interneurons) and neck muscles (via motor neurons in S2). Shallow papillary circuits are associated with deterministic behaviors, while amphid sensory responses are associated with probabilistic/plastic behaviors^35,49,50,54^. Circuits are associated with deterministic and probabilistic behaviors, with individual neuron classes and known connectivity, schematized from sensory input to motor outputs. Individual neuron classes and muscle outputs are colored according to the strata they belong to (for muscles, according to the strata innervating neurons belong to). **d-e,** Axial view of the ‘rich-club’ AIB left (AIBL) interneuron^18,21^ (**Fig. 2f**) with distribution of the presynaptic (**d**) and postsynaptic (**e**) sites colored according to the strata of the corresponding AIBL synaptic partner. Vertical dashed line indicates division between proximal and distal parts of the neurite. Similar distribution was seen for AIB right (AIBR, not shown). **f,** Volumetric rendering of ‘rich-club’ interneuron RIA right (RIAR). Interneuron RIA innervates two strata, amphid-sensory rich S3/S4 with motor-neuron rich S1/S2. Similar distribution was seen for RIA left (RIAL, not shown). The morphology of RIA in the context of the strata, and the polarized distribution of its synapses, are consistent with its known role from behavioral studies of transducing amphid-sensory information to modulate head movements^83–86^ (circuit schematic in **(c)**).

**Extended Data Figure 3. S1 honeycomb structure precisely encases the S2, S3, and S4 bundles of the neuropil**

**a,** Illustration of the IL1L neuron adapted from^3^. Schematic of worm head and neuropil (dashed box) above. In schematic below, IL1 neuron in red, with dendrite (left pointing arrow), soma (circle) and axon (right pointing loop, arrow). Pharynx (shaded gray) for reference. **b**, Cartoon of IL1L loop in the context of the nerve ring neuropil (sensory dendrite not show). **c**, Volumetric reconstruction of IL1L from L4 animal EMs. **d**, Overlay of IL1L volumetric reconstruction and synapses (grey squares) highlighting the position of the synaptic endplate (after the neuron loops around the neuropil). **e,** Volumetric reconstruction of the L4 *C. elegans* neuropil (from EM serial sections^3^) with individual neurons from the 4 strata colored (S1-Red, S2-Purple, S3-Blue, S4-Green). Scale bar is 5 µm. **f-g**, Schematic of the scaffold formed by the S1 looping structures, from a lateral (**f**) and axial (**g**) view with neuropil strata. **h**, Schematic of the S1 loops (lateral view) without the strata (as **Fig. 1d, e**) **i-j,** Volumetric reconstruction of Dorsal Right Loop and S2 (**i**) and S3/S4 (**j**) with individual neurons that fall outside of the looping structure rule (yellow arrows). Although 90% of S2, 84% of S3 and 100% of S4 neurons are contained within the indicated loops, a minority of neurons belonging to S2 and S3 strata are not encased by the loops corresponding to the specific strata. We include the names of these neurons below the volumetric reconstructions; Supplemental Video 5. **k,** List of all neurons and their positions in the dorsal right loop structures, colored according to the 4 strata (a complete listing of all neurons, and their positions within the 6-fold symmetric honeycomb scaffolding structure, can be found in Supplemental Table 1).

**Extended Data Figure 4. Neurons unassigned to the 4 strata anatomically contact multiple strata and belong to the highly interconnected “rich-club” neurons.**

**a-d,** Volumetric reconstructions of the unassigned AVE interneuron (yellow) in the context of nerve ring strata, with AIB (grey). Arrows indicate the two segments of AVE that border strata. AVE has a similar morphology to AIB (**Fig. 2f**) but is anteriorly displaced by one stratum: AVE borders S2/S3 (**b-c**), shifts along the A-P axis, and then borders S1/S2 (**c-d**). Lines in (**a,b**) indicates AVE shift along the A-P axis to shift strata; Supplemental Video 7. **e,** Analysis of the total number of neurons within each stratum, and in the unassigned group. **f,** Classification of ‘unassigned’ neurons. **g,** Stratum location of the rich club neurons^18,21^. Colored box depicts strata assignment. **h-u,** Volumetric reconstructions of all unassigned neurons highlighting their strata interactions. **h,** AIZ shifts perpendicularly from S3 to the S2/S3 border, highlighted with arrow. **i,** RIB forms a cage-like structure around S2. **j,** AVA borders S2/S3 and protrudes into S3 and S4. **k,** PVR borders S1/S3 and protrudes into S1. **l,** RIM borders S2/S3 and protrudes into S3. **m,** RMG protrudes into S1/S2/S3. **n,** SDQ borders S2/S3. **o,** SIBV’s main neurite is in S2, but it sends a second neurite into S1. **p,** URX interacts with S1 and S4. **q,** AWA borders S3/S4. **r,** RIG borders S2/S3. **s,** RIR borders S3/S4. **t,** RIS borders S1/S2. **u,** FLP has sparse segmentation data in the nerve ring. Images are rotated relative to each other, and transparency setting vary between images, for clarity in display of their position within the nerve ring.

**Extended Data Figure 5. Analysis of the 4 strata in the context of the lineage tree**

**a,** Lineage tree for *C. elegans* (0-428 minutes post fertilization). Each neurons’ terminal branch is colored according to its stratum assignment. Upon detailed examination of all 181 neuropil neurons in the context of the lineage tree, we could not identify correlations between terminal lineage position and stratum assignment.

**Extended Data Figure 6. Early segregation of neuronal somas followed by S2 pioneering neuron outgrowth initiate development of the stratified neuropil**

**a-d,** Neuronal soma positions within the embryo (generated using WormGUIDES^66^). Somas are colored according to their future neurite strata assignment. S2 pioneer neuron, SIAD, in white for reference of nerve ring position. White arrowheads in (**c,d**) highlight the growing tips of the pioneer SIAD. White arrowhead in (**e**) highlights the dorsal midline (meeting point for the bilateral SIADs). Note that S1 somas are anteriorly segregated prior to pioneer neurite outgrowth. Supplemental Video 8. **f-h,** 3D depth trajectories displaying the movement of cells which will extend their neurites to S2 (**f**), S3 (**g**) or S4 (**h**) and colored accordingly (movement represented for timepoints 380-420 mpf). S2 movement has a ventral bias. S3 movement is principally along the outermost embryonic edge, and S4 clusters into 2 bilaterally symmetric groups. **i,** Schematic of *C. elegans* embryo depicting region displayed in (**j-n**), red box. **j-n**, Time-lapse of the outgrowth dynamics of the neuropil (labeled with ubiquitous Pnhr-2::membrane-tethered::GFP) and schematic. Arrowheads indicate first extensions entering the future neuropil. Deconvolved diSPIM maximum intensity projections are shown; same dataset as Supplemental Video 9. **o,** Schematic of *C. elegans* embryo depicting region displayed in (**p-r**), red box. **p-r** Comma stage embryo (400 min) co-labeled with pan-neuronal (Pnhr-2::membrane-tethered::GFP) and pioneer neuron (Plim-4::mCherry) markers. Membrane and pioneer expression co-localize in SIAD, SIBV, and SMDD confirming the lineaging analysis (see Fig. 3d-g). Single z-slice acquired with one diSPIM arm shown. Arrow indicates co-labeling of the two markers. **s,** Schematic of *C. elegans* embryo depicting region displayed in (**t-v,w-y**), red box. **t-v**, Time-lapse of the outgrowth dynamics of pioneer neurons (labeled with Plim-4::membrane-tethered::GFP). The same images were used in Fig. 3h and displayed here for comparison with the next panels. Deconvolved diSPIM maximum intensity projections are shown. **w-y**, Time-lapse of the outgrowth dynamics of the neuropil (labeled with Prab-3::membrane-tethered::GFP). Deconvolved diSPIM maximum intensity projections are shown. Neuronal outgrowth into the neuropil occur simultaneously for pioneers (**t-v**) and the first pan-neuronal outgrowth events detected (**x-y**), suggesting the S2 pioneers are the first to enter the developing neuropil; same dataset as Supplemental Video 11. **z,** Analysis of timing for arrival to the dorsal midline in a strain co-labeled with Prab-3::membrane-tethered::GFP, and Plim-4::mCherry. Points connected with a line correspond to two datapoints from the same animal (n=10). “ns” not significant by paired Student’s t test. Scale bar (10 µm) applies to panels in corresponding sections (**j-m, p-r, t-v, w-y**). Timing for all panels is mpf.

**Extended Data Figure 7. S2 Pioneer neurons are required for nerve ring neuropil development.**

**a,** Schematic of caspase ablation strategy for pioneer neurons. Top panel depicts split-caspase induced cell ablation, as described in^77^. Bottom panel depicts pioneer specific split-caspase ablation assay in embryos. **b-c,** Time-lapse of the outgrowth dynamics of the pioneering neurons in control (**b**) and pioneer ablated (**c**) embryos (labeled with Plim-4::mCherry). Pixel intensities are different for (**b,c**) due to significant decrease in signal in the ablated animals. Deconvolved diSPIM maximum intensity projections are shown. Scale bar (10 µm) applies to all panels. **d,** Quantification of the percentage of embryos forming a full neuropil ring in control and pioneer ablation embryos. ‘n’ corresponds to the number of embryos scored. ****p<0.0001 by Fisher’s exact test between Ctrl and Ablation. **e,** Time-lapse of the dynamics of 2xFYVE on ablated pioneering neuron somas. 2xFYVE is a marker of cell death and appears around cell corpses as described^80^. To see cell corpses of pioneer neurons, embryos were labeled with pced-1_2xFYVE_GFP(S65C,Q80R) (to image cell corpses) and Plim-4::mCherry (to image pioneer neurons). Single Z-plane from diSPIM dataset shown. Scale bar (3 µm) applies to all panels. **f,** Quantification of 2xFYVE encasing ablated pioneer somas. **p<0.01 by Fisher’s exact test between Ctrl and Ablation. **g,** Volumetric reconstruction of the developing neuropil for control and pioneer ablated embryos. Volumes were acquired from diSPIM images analyzed with 3D Object Counter (FIJI-ImageJ2; see Methods). Green arrowheads emphasize aberrant neuropil phenotypes in ablation animals (gaps in the neuropil and decreased widths). **h,** Analysis of pixel intensity within the neuropil volume of control and pioneer ablated embryos. Each dot represents the summation of all pixels within a neuropil volume for 1 embryo quantified using 3D Object Counter (FIJI-ImageJ2 and see Methods). **p<0.01 by Student’s t test between Ctrl and Ablation. Error bars are SEM. Timing for all panels is mpf.

**Extended Data Figure 8. S2 Pioneer neurons are required for the development of neurons from all 4 strata, and the unassigned neurons.**

**a,** Schematic of embryo highlighting area in (**b-c**) as red rectangle. **b-c,** Time-lapse of outgrowth dynamics of Stratum 3 neuron ASH in control (**b**) and pioneer ablated (**c**) embryos, with schematic (right hand side). Deconvolved diSPIM maximum intensity projections shown; images in (**b**) same dataset as Supplemental Video 14. **d-e,** Quantifications of ASH axon (**d**) or dendrite (**e**) outgrowth for control and ablated animals. Note how axons (which are in the nerve ring) are affected by nerve ring pioneer neuron ablations, while dendrites (which are not in the nerve ring) are not affected. ‘n’ = number of neurons quantified. *p<0.05, **p<0.01, ***p<0.001, ****p<0.0001 by unpaired Student’s t test between Ctrl and Ablation at each timepoint. Timepoints without annotation are “ns” not significant. Error bars are SEM. **f-i**, As for (**a-d**), but for unassigned interneurons AIB; images in (**g**) same dataset as Supplemental Video 15. **j-n,** As for (**a-e**), but for Stratum 4 neuron BAG. **o**, Quantification of the percentage of BAG neurons with defective morphologies at 444 mpf for control and pioneer ablated animals. Note that BAG neurons are delayed in early outgrowth, but eventually find their terminal locations, suggesting guidance of this neuron relies on redundant mechanisms. ‘n’ = number of embryos. “ns” not significant by Fisher’s exact test between Ctrl and Ablation. **p,** As for (**d**) but for Stratum 2 neuron AVA shown in (**Fig. 4d,e**). **q,** Schematic of *C. elegans* head highlighting area in (**r-s**) as red rectangle. **r-s**, Larval stage 1 (L1) images of Stratum 1 neuron OLL in control and pioneer ablated animals, with schematic (below). L1 images were taken because there were no available promoters to image OLL in embryos. Spinning disk confocal maximum intensity projections shown. **t-v,** As for (**q-s**) but for Stratum 3 neuron AIY. **w-y,** As for (**q-s**) but for unassigned neuron AIB. Note that AIB outgrowth defect in embryogenesis (**h**) persists to larva stage 1 (L1) (**y**). **z,** Quantification of the percentage of AIB neurons with defective morphologies in L1 animals for control and pioneer ablated animals. ‘n’ = number of animals scored. ***p<0.001 by Fisher’s exact test between Ctrl and Ablation. For cell specific labeling of neurons, see Methods and Supplemental Tables 2 and 3. Scale bar (10 µm) applies to panels (**b-c,g-h,k-l,r-s,u-v,x-y**), and timing for all panels is mpf. Neurons are colored according to which strata they belong (S1-Red, S2-Purple, S3-Blue, S4-Green, Unassigned-Yellow).

**Extended Data Figure 9. A temporal progression of outgrowth, beginning with the S2 Pioneers, results in the inside-out development of the nerve ring.**

**a-d,** Time-lapse of the outgrowth dynamics of S3 neuron ASH in control animal. For ASH cell-specific labeling see Methods. **a’-d’,** As (**a-d**) but includes S2 pioneer neurons (labeled with Plim-4::mCherry). Yellow arrowheads marks ASH axonal outgrowth in the context of the pioneers. White arrowhead marks dorsal midline. ASH outgrowth into the neuropil occurs after the pioneers have grown into the nerve ring. Deconvolved diSPIM maximum intensity projections shown. **e-i,** Time-lapse of the outgrowth dynamics of unassigned neuron AIB in control animal. For AIB cell-specific labeling see Methods. **e’-i’,** As (**e-i**) but includes S2 pioneer neurons (labeled with Plim-4::mCherry). Blue arrowheads marks AIB axonal outgrowth in the context of the pioneers. White arrowhead marks dorsal midline. Note that AIB enters the neuropil after the pioneers and as ASH has reached the dorsal apex (compare **d’** to **g’**). Data collected in this way (**a-i**) were used for indicated neurons in **Fig. 4j**. Deconvolved diSPIM maximum intensity projections shown. **j-q,** Time-lapse of the outgrowth dynamics for S2 pioneer SAAV, and S4 neurons (AFD, ADF, AWC, AWB). In (**j**) red arrowheads mark outgrowth of SAAV. In (**k-q**) red arrowhead marks dorsal midline and yellow arrowhead marks outgrowth of S4 neurons. The S4 neurons pause for ∼30 min near the SAAV soma before growing into the nerve ring. Deconvolved diSPIM maximum intensity projections shown. Scale bar (10 µm) applies to panels (**a-d’, e-i’, j-q**), and timing for all panels is mpf.

## Supplemental Videos

**Supplemental Video 1. C-PHATE plot reveals hierarchical nested clustering over the DC iterations for L4 *C. elegans*.**

3D rendering of C-PHATE plot of DC analyses for a larva stage 4 (L4) animal (DC iterations 1-11 shown). Individual neurons are located at edges of the graph and condense as they move centrally. Plot colored according to 4 super-clusters (C1-Red, C2-Purple, C3-Blue, C4-Green).

**Supplemental Video 2. C-PHATE plot reveals hierarchical nested clustering over the DC iterations for adult *C. elegans*.**

3D rendering of C-PHATE plot of DC analysis for an adult stage animal (DC iterations 1-10 shown). Individual neurons are located at edges of the graph, with condensation iterations towards the centerPlot colored according to 4 super-clusters (C1-Red, C2-Purple, C3-Blue, C4-Green). Unlike JSH, N2U has two smaller clusters that are not part of the 4 super clusters, and these are colored in grey (see Methods for neuronal classification parameters for bundle identification).

**Supplemental Video 3. The nerve ring neuropil is structurally organized into 4 distinct stratified bundles.**

Volumetric reconstruction of the L4 *C. elegans* neuropil (from EM serial sections^3^) with neurons from the 4 strata highlighted (S1-Red, S2-Purple, S3-Blue, S4-Green).

**Supplemental Video 4. S1 neurons form a honeycomb-like scaffold encasing S2, S3, and S4.**

Volumetric reconstruction of the L4 animal honeycomb-like scaffold formed by the 32 neurons in S1. During first part of the movie, bilaterally symmetric neuron pairs are highlighted in a range of colors as to differentiate them. In the second part, the color scheme changes, with the entire scaffold shown in red (due to their placement in S1) in the context of the remaining strata (S2-Purple, S3-Blue, S4-Green).

**Supplemental Video 5. The S1 honeycomb-like scaffold precisely loops at the boundaries of the indicated strata.**

Volumetric reconstruction of a subset (dorsal right) loop neurons forming the honeycomb scaffold around the strata (Neurons: URADR, URYDR, IL1DR, and IL2DR, in red), in the context of S2 bundle (purple), and then S3/S4 bundle (blue/green).

**Supplemental Video 6. Unassigned interneuron AIB contacts multiple strata**

Volumetric reconstruction of the unassigned (yellow), ‘rich-club’ AIB interneurons^18,21^ in the context of nerve ring strata. AIB borders S3/S4, shifts along the A-P axis, and then borders S2/S3.

**Supplemental Video 7. Unassigned interneuron AVE contacts multiple strata**

Volumetric reconstructions of the unassigned (yellow), ‘rich-club’ AVE interneuron^18,21^ in the context of nerve ring strata and AIB (orange). AVE borders S2/S3, shifts along the A-P axis, and then borders S1/S2.

**Supplemental Video 8. Early segregation of neuronal somas followed by S2 pioneering neuron outgrowth initiate development of the stratified neuropil**

Representation of the neuronal soma positions within the embryo and S2 pioneer SIAD outgrowth between 330-427 mpf. Data were generated using the WormGUIDES atlas^66^. Briefly, record of the position of every cell at every minute during embryonic development for three embryos were used to generate a 4D representation of all cellular positions as described^66^. To generate the movie, these datasets were inspected with somas colored to show their eventual neurite strata assignment (S1-Red, S2-Purple, S3-Blue, S4-Green), and pioneer SIAD neurons in white to pinpoint the position of the developing neuropil. Note that S1 is anteriorly segregated prior to pioneer neurite outgrowth.

**Supplemental Video 9. Embryonic neuropil development begins between 390-400 mpf.**

Time-lapse of the outgrowth dynamics of the neuropil (labeled with ubiquitous Pnhr-2::membrane-tethered::GFP). Neuropil visible as “horseshoe” structure in the embryo head (anterior of the embryo is towards the top). Images are deconvolved diSPIM maximum intensity projections. Time-lapse images in Extended Data Fig. 6j-m were generated from this dataset. Scale bar applies to entire movie, and timing is mpf.

**Supplemental Video 10. S2 pioneering neurons bundle together and enter the presumptive neuropil between 390-400 mpf.**

Time-lapse of the outgrowth dynamics of pioneer neurons (labeled with Plim-4::membrane-tethered::GFP, followed by a 3D rotation of the last timepoint to highlight the early ring structure formed by the S2 pioneers. Images are deconvolved diSPIM maximum intensity projections. Time-lapse images in Fig. 3h-i are from this dataset. Scale bar applies to entire movie, and timing is mpf.

**Supplemental Video 11. Embryonic neuropil development begins between 390-400 mpf.**

Time-lapse of the outgrowth dynamics of the nerve ring (labeled with Prab-3::membrane-tethered::GFP), followed by a 3D rotation of the last timepoint to highlight the neuropil, which is the bright ring structure in the anterior part of the embryo (top). Embryo starts twitching during later development. Note that the neuropil increases in intensity overtime, probably due to an increase in the number of neurons entering the neuropil through time. Images are deconvolved diSPIM maximum intensity projections. Time-lapse images in Extended Data Fig. 6w-y are from this dataset. Scale bar applies to entire movie, and timing is mpf.

**Supplemental Video 12. Embryonic neuropil development begins between 390-400 mpf.**

Time-lapse of the outgrowth dynamics of the nerve ring (labeled with ceh-48::membrane-tethered::GFP), followed by a 3D rotation of the last timepoint to highlight the neuropil, which is the bright ring structure found near the top of the embryo. Embryo starts twitching during later development. Note that the neuropil increases in intensity overtime, probably due to an increase in the number of neurons entering the neuropil through time. Images are deconvolved diSPIM maximum intensity projections. Scale bar applies to entire movie, and timing is mpf.

**Supplemental Video 13. Outgrowth dynamics of the S2 neuron AVL.**

Time-lapse of the outgrowth dynamics of the AVL neuron (labeled with lim-6::GFP), followed by a 3D rotation of the last timepoint to highlight AVL outgrowth into the neuropil. Embryo starts twitching during later development. Images are deconvolved diSPIM maximum intensity projections. Time-lapse image in Fig. 4d is from this same dataset. Scale bar applies to entire movie, and timing is mpf.

**Supplemental Video 14. Outgrowth dynamics of the S3 neuron ASH.**

Time-lapse of the outgrowth dynamics of the ASH neuron (labeled with Punc-42::ZF1::membrane-tethered::GFP + Plim-4::ZIF-1, see Methods, followed by a 3D rotation of the last timepoint to highlight ASH outgrowth into the neuropil. ASH is the brightest neuron seen in the movie. Embryo starts twitching during later development. Images are deconvolved diSPIM maximum intensity projections. Time-lapse images in Extended Data Fig. 8b are from this dataset. Scale bar applies to entire movie, and timing is mpf.

**Supplemental Video 15. Outgrowth dynamics of the unassigned neuron AIB.**

Time-lapse of the outgrowth dynamics of the AIB neuron (labeled with Punc-42::ZF1::membrane-tethered::GFP + Plim-4::ZIF-1, see Methods), followed by a 3D rotation of the last timepoint to highlight AIB outgrowth into the neuropil. AIB is the brightest neuron seen in the later time points. Images are deconvolved diSPIM maximum intensity projections. Time-lapse images in Extended Data Fig. 8g are from this same dataset. Scale bar applies to entire movie, and timing is mpf.

## Materials and Methods

### Diffusion condensation (DC) coarse graining of electron micrographs of the nerve ring connectome

We used segmented data of all neurons for the two available connectomes^3^ and performed adjacency analyses as described^22^, with the following modifications: we excluded segmented cell bodies (identified by the dark staining of the nucleus) and analyzed the following EM slices corresponding to the nerve ring: (1-282) for JSH dataset and (2-182, and VC1-34) for N2U datasets (available at www.wormwiring.org^22^). This algorithm extracts: 1) neuron-neuron adjacencies and 2) the number of pixels each adjacency persists for (termed the “weight” of the adjacency).

We developed a novel algorithm called Diffusion Condensation (DC^7^) to analyze the two neuronal adjacency datasets. We noted that neurons have several levels of hierarchical organization, from single neurons to circuits to bundles. Therefore, rather than choosing a clustering algorithm which provides one clustering scale, we choose an algorithm that reveals structure at all scales, uncovering both fine circuits and also gross nerve ring architecture.

Before running DC, an *n* by *n* adjacency matrix was built by summing the weight of each neuron-neuron contact over all EM slices. This square and symmetric matrix, where each row and each column represent the connection profile of that neuron with every other neuron in the nerve ring, is row normalized such that each row sums to one, creating a Markov matrix. This normalized matrix is then eigendecomposed and the first 50 eigenvectors extracted, creating a low dimensional embedding of the neurons’ connectivity profile.

DC leverages this low dimensional embedding of the neurons to capture the intrinsic manifold geometry by using random walks that aggregate local affinity to reveal nonlinear relations in the data. These local affinities are constructed using a gaussian kernel:

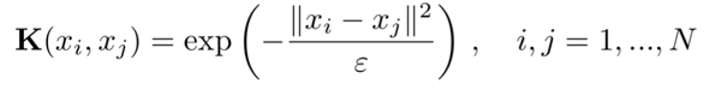

Where K is an *n* by *n* square matrix and (i, j) denote individual neurons. Here the bandwidth parameter ε controls neighborhood size. DC first constructs affinities with a small bandwidth, only allowing neurons that are very similar to one another be connected in matrix K. Using the affinity matrix K, a diffusion operator P is created where **P** = **D^−1^K** and D is the diagonalized row sum of K. The diffusion operator P defines a set of transition probabilities of a random walk over the eigenvectors of the adjacency matrix. By apply P to our eigenvectors, **X’** = **X P**, DC then moves points in our input matrix, the eigenvectors of the adjacency, to local centers of gravity, reducing variability between similar points.

As the DC process continues, transition probabilities are iteratively calculated with increasingly large bandwidths, and applied to the output of the previous DC run. Such an iterative process diffuses the data with larger and large bandwidths, connecting points that are more dissimilar and moving them towards a shared center of gravity. The effect this has, in practice, is to first remove local variability between similar neuron’s connection profiles, before scaling to remove differences in more dissimilar neuron connection profiles. When neurons’ connection profiles become similar and the distance between these points falls below a certain preset threshold *t*, neurons are grouped together as being part of the same cluster. This strategy allows for groupings of similar neurons at all granularities. In this way, DC creates a hierarchical tree of neuron contact similarities, revealing small circuits of neurons at fine grain and large bundles at coarser grain.

DC was performed on a computer with the Windows7 operating system (Processor Intel(R) Core(TM) i7-6700K CPU @ 4.00GHz, 4008 Mhz, 4 Core(s), 8 Logical Processor(s)), Memory: 16GB) using MATLAB R2017b. The DC algorithm is a deterministic algorithm and produces the same result on every run. However, results may slightly vary due to the parameter t (which stores the affinity threshold at which points are merged to create groupings). Of note, computers with different machine precisions can store t with different numbers of bits, causing the algorithm to vary slightly in the number of iterations needed until the entire dataset merges. Code for DC can be found on https://github.com/agonopol/worm_brain.

After DC analysis, we calculated network modularity scores for each cluster within each iteration as described^41^. Adjusted rand index (ARI) was computed at each condensation iteration between N2U and JSH as described^87^, and the mean ARI reported as ARI = 0.7 (for −1 ≤ ARI ≤ 1).

We inspected the results for the DC analysis, focusing on the iterations that had the highest modularity score: iteration 23 for JSH and 22 for N2U, which showed the four reported super-clusters. We inspected cell morphologies for each of the individual neurons belonging to each stratum, and correlated their identities, morphologies and position in the DC analyses, for both JSH and N2U datasets. Because most neurons are bilaterally symmetric, this meant we inspected, for most neuron classes, placement of four individual neurons within each stratum (two bilaterally symmetric individual cells for the JSH dataset, and two for the N2U dataset). We adopted a threshold criterion for accepting inclusion of neurites in the consensus cell lists for the four strata: they were assigned to a stratum if in 3 of the 4 analyzed instances the individual neurons were placed in the same cluster by the DC analyses. For neurons that were not bilaterally symmetric, correspondence of the single neurons between N2U and JSH was used to assign strata. Neurons that fell beneath this threshold were catalogued as “unassigned” (**Extended Data Fig. 1)**.

### C-PHATE visualization of electron micrographs of the nerve ring connectome

To visualize the DC tree we adapted M-PHATE and PHATE^30,88^ to generate C-PHATE (for Condensation PHATE). At each DC time point, we extracted the intra-timepoint kernel K, an n x n matrix that denotes affinities between each of n points. To capture the evolution of these intra-timepoint kernels over DC, we co-embed them in a multi-timepoint kernel, where a kernel from each timepoint was placed along the diagonal. Similar to M-PHATE^88^, in order to connect the kernels together in our tree embedding, points from successive kernels were connected with a weight. Different from M-PHATE, however, points that are assigned to the same cluster in successive iterations are also connected with a weight in the multi-timepoint kernel. This multi-timepoint kernel is plotted via the PHATE algorithm^30^ to reveal the hierarchical structure created by DC and to visualize both the dynamics of the condensation process as well as the relationship to one another at all levels of the hierarchy.

### Renderings of neurites and synapses in the EM datasets

Segmentations of neuron membrane boundaries from the EM datasets^22^ were rendered in 2D using TrakEM2 in Fiji (ImageJ2)^89^. Segmentations in TrakEM2 were then rendered volumetrically by using the 3D viewer plugin preinstalled in Fiji (ImageJ2; downloaded from https://imagej.net/Fiji#Downloads) and saved as object files (.obj). Renderings were then generated in 3D viewer Fiji (ImageJ2) or Blender (downloaded from https://www.blender.org/download/). CytoSHOW, an open source image analysis software^62^, was used to generate the C-PHATE 3D renderings by marking the coordinates of each cluster-iteration-point with an info-linked icosphere, color coded by stratum, and saved as an obj file. CytoSHOW can be downloaded from http://www.cytoshow.org/ as described^62^.

Supplemental Video 3, which integrates volumetric segmentation and the electron microscopy images, was generated by exporting all segmentations within TrakEM2 as labels, and EM images as tiffs. All images and labels were identically resampled and saved as tiff stacks, which were imported into Microscopy Image Browser version 2.511 (MIB)^90^. Images were then exported as .mrc files and models as .mod files, and opened in 3dmod (IMOD)^91^.

Synapse position data (**Extended Data Fig. 2d,e and 3d**) in the context of the volumetric renderings were generated by importing from the WormWiring project’s Elegance format into CytoSHOW for display as info-linked tags overlaid on the segmented EM sections^11,22^. The same unwarped registered serial section image stacks were used in both the synaptic and cell-shape cataloging projects by the Emmons group^11,22^. We therefore imported cell-outline data from TrakEM2 format into CytoSHOW as info-linked overlay tags. By making minor corrective shifts through visual inspection in the x, y, and z synapse coordinates, we achieved nanometer-precision alignment of neurite outlines with synapse markers within an integrated CytoSHOW tag set.

### Lineage and soma movement analyses via WormGUIDES atlas

Using the WormGUIDES atlas^66^ available online (https://wormguides.org/wormguides-atlas/), we colored all somas according to their eventual neurite stratum assignment. Note that the timing reference is different between our analysis and the WormGUIDES atlas (our timing is referenced to minutes post fertilization, and the WormGUIDES atlas is referenced to minutes post first cell cleavage). For lineage analysis we used the Lineage Tree function built into the WormGUIDES atlas. To analyze soma movements, we tracked all the neuronal somas over time within the WormGUIDES atlas. 3D depth trajectories displaying movement of cells for the different strata were generated in MATLAB as described^92^. Cell motion was summarized with a smooth truncated cone connecting the cell’s start and end position over the period analyzed.

### *C. elegans* strains

*C. elegans* strains (genotypes in Supplemental Table 2) were raised at 20°C using OP50 *Escherichia coli* seeded on NGM plates. The wild-type reference strain was N2 Bristol. We received strain OH812 and OH1422 from the Caenorhabditis Genetics Center (CGC).

### Molecular biology and Transgenic lines

Plasmids used to generate transgenic *C. elegans* strains were made by Gateway system (Invitrogen) or Gibson Assembly system (New England Biolabs). For a list of plasmids and the primers used see Supplemental Table 3 Detailed cloning information will be provided upon request. Transgenic strains injected with (10-100 ng/uL) of plasmid DNA were generated via standard techniques,^93^ with larva co-injection markers Punc-122::GFP or Punc-122::RFP, or embryonic co-injection markers Pelt-7::mCherry::NLS or Pelt-7::GFP::NLS. A plasmid with Pceh-48 was provided by Oliver Hobert (Columbia University, New York City, New York). A plasmid with 2xFYVE was provided by Zheng Zhou^80^ (Baylor College of Medicine, Houston, Texas).

### Imaging, Lineaging and quantification of outgrowth

Larval stage 1 animals (L1) were imaged in a UltraView VoX spinning-disc confocal microscope (PerkinElmer Life and Analytical Sciences). Embryonic imaging was performed via dual-view inverted light sheet microscopy (diSPIM)^64,65^. For all embryonic imaging, onset of twitching was used to time embryonic development. Onset of embryonic twitching is stereotyped and starts at 430 minutes post fertilization (mpf) for our imaging conditions.

Lineaging was performed using StarryNite/AceTree^63,73,74^, and the approaches were integrated as described^62^. Lineaging information for specific promoters is available at http://promoters.wormguides.org. For lineaging of the pioneer neurons, three embryos were lineaged to assess reproducibility of lineaged identities.

Imaging of outgrowth dynamics (**Extended Data Fig. 9j-q**) was performed using diSPIM and a Yokogawa CSU-X1 spinning-disk unit. Outgrowth of neurites was monitored by scoring time to arrival at the dorsal midline. RMDD analysis (**Fig. 4j**) was performed on datasets from^62^. Pan-neuronal and pioneer growth to dorsal midline (**Extended Data Fig. 6z**) were scored blindly to genotypes for animals co-expressing both markers.

### Pioneer Neuron Ablation Assay

We used caspase ablation assays^77^ to ablate pioneer neurons. Integrated *C. elegans* lines expressing either the p12 or p17 subunit of human Caspase-3 driven by the lim-4 promoter were independently generated (see Supplemental Tables 2 and 3 for strain and plasmid information). Strains used were DCR6102 and DCR6100.

To induce caspase ablations, DCR6102 was crossed to DCR6100, and comma stage F1 embryos were imaged on the diSPIM. Neurite outgrowth for all experiments was assessed by co-injecting with transgenes of interest and an embryonic co-injection marker (Pelt-7::mCherry::NLS), or by crossing to integrated neuronal markers (see Supplemental Table 2 for details). No pioneer neurite outgrowth was seen after pioneer caspase ablations. Cell death scoring was performed by monitoring 2xFYVE during pioneer ablations as described^80^. Controls correspond to array-expressing N2 animals without the caspases.

Quantification of neuropil volume and volumetric intensity during pioneer ablations was performed using 3D Objects Counter in Fiji (ImageJ2). Single view diSPIM images of embryos expressing Pceh-48::membrane-tethered::GPF were analyzed. The thresholds used in 3D Objects Counter were calculated by quantifying the ‘background’ fluorescent intensity in tail region (where there is no neuropil signal but there is background tissue signal), and then normalizing to the highest recorded background intensity. The thresholds were used in 3D Objects Counter to volumetrically segment the nerve ring in each embryo.

Quantification of neuronal morphologies following pioneer ablations (**Fig. 4f**) was performed by scoring neuronal morphology at the time when control animals’ neurons had reached the dorsal midline. When the neuron scored was a bilateral pair, the embryo/animal was considered ‘abnormal’ if either neuron’s morphology was abnormal. AVL, AIB, ASH, and BAG were scored in embryos, and OLL, AIY, and AIB were scored in L1 animals (**Fig. 4f and Extended Data Fig. 8**).

Quantifications of neuronal outgrowth in control and pioneer ablated animals were analyzed using Simple Neurite Tracer v2.0.2 in Fiji (ImageJ2). Deconvolved diSPIM images were used to trace the neurites from the soma to the neurite tip. Each timepoint shown is representative of at least 3 neurons scored. Same datasets were used to quantify and compare aberrant outgrowth, dorsal midline timing, and outgrowth dynamics for **Fig. 4d,e,f,j** and **Extended Data Fig. 8**.

### AIB and ASH imaging

Lineaging analysis determined that the unc-42 promoter labels AIB, ASH, and the pioneering neurons during embryogenesis. To generate a strain that allowed for individual AIB or ASH analyses we applied the ZIF-1/ZF1 system as described^94^. Briefly, we generated a strain containing Punc-42::ZF1::Membrane-tethered::GFP and Plim-4::SL2::ZIF-1, which degraded GFP in the pioneering neurons. In the pioneer ablation experiments in which we analyzed AIB and ASH, only the construct Punc-42::Membrane-tethered::GFP was used.

### Statistical analysis

Specified statistical analyses for continuous data were based on student’s t-test for comparisons between two groups or one-way ANOVA with post hoc analysis by Turkey’s multiple comparison test for three or more groups. Error bars were reported as standard errors of the mean (SEM). For categorical data, groups were compared with Fisher’s exact test. All were analyzed using Prism 8.

