## Extended Data Figure 1 for "Structural and developmental principles of neuropil assembly in *C. elegans*"

Extended Figure 1: Moyle et al.

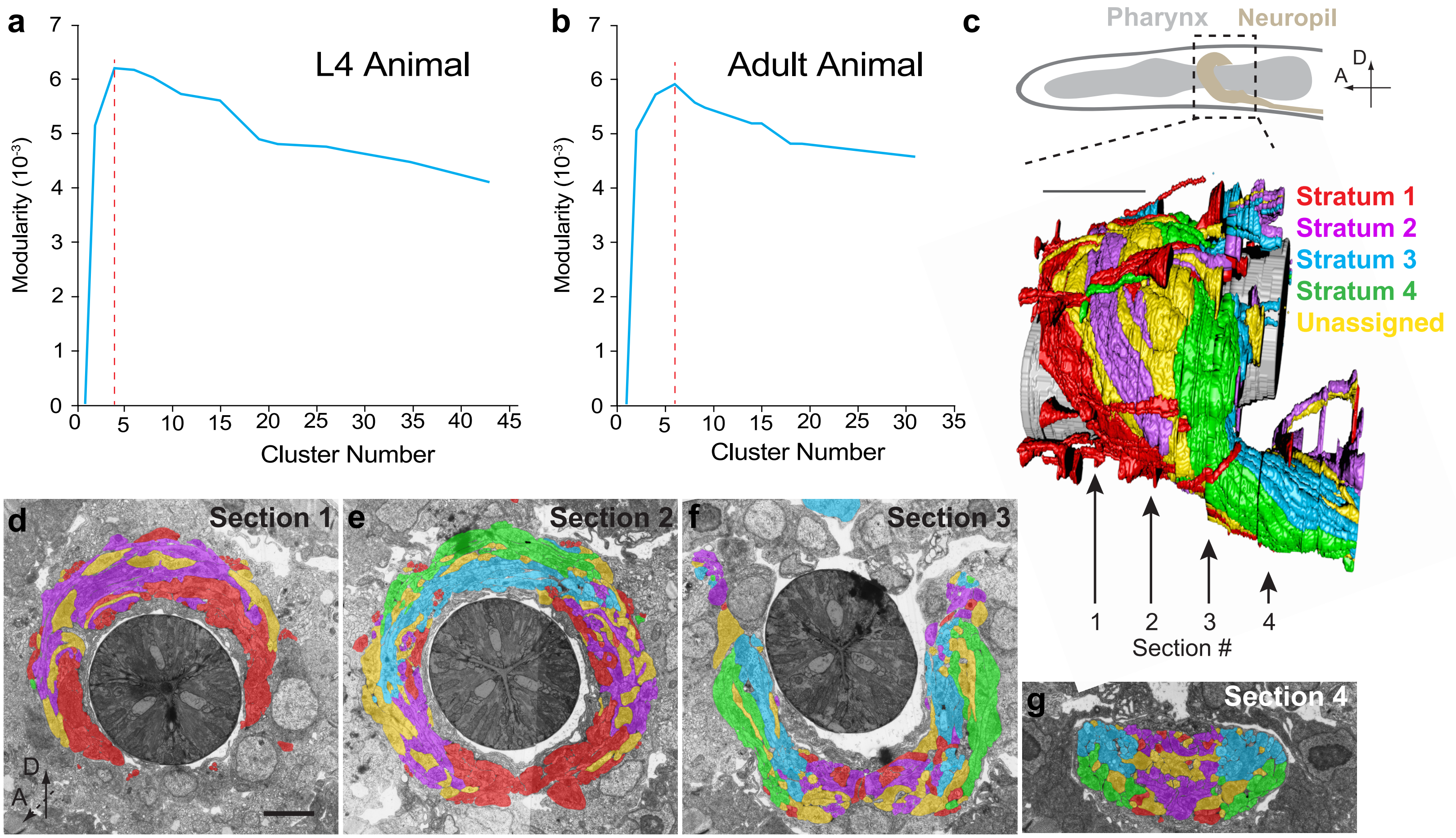


**h**

| Stratum 1 | Stratum 2 | Stratum 3 | Stratum 4 | Unassigned |
|---|---|---|---|---|
| Total: 45 | Total: 39 | Total: 43 | Total: 24 | Total: 30 |
| CEP (4) | ADE (2) | ADA (2) | ADF (2) | AIB (2) |
| IL1 (6) | ALN (2) | ADL (2) | AFD (2) | AIZ (2) |
| IL2 (6) | AVK (2) | AIM (2) | AIA (2) | AVA (2) |
| OLL (2) | AVL (1) | ALA (1) | AIN (2) | AVE (2) |
| OLQ (4) | DVC (1) | ALM (2) | AIY (2) | AWA (2) |
| RIA (2) | PLN (2) | AQR (1) | ASE (2) | FLP (2) |
| RIH (1) | PVT (1) | ASH (2) | ASG (2) | PVR (1) |
| RIP (2) | RIC (2) | ASJ (2) | ASI (2) | RIB (2) |
| RMDD (2) | RIV (2) | ASK (2) | AUA (2) | RIG (2) |
| RMDV (2) | RMD (2) | AVB (2) | AWB (2) | RIM (2) |
| RME (4) | RMF (2) | AVD (2) | AWC (2) | RIR (1) |
| URA (4) | RMH (2) | AVF (2) | BAG (2) | RIS (1) |
| URB (2) | SAA (4) | AVH (2) | | RMG (2) |
| URY (4) | SIA (4) | AVJ (2) | | SDQ (2) |
| | SIBD (2) | AVM (1) | | SIBV (2) |
| | SMB (4) | BDU (2) | | URX (2) |
| | SMD (4) | DVA (1) | | VB01 (1) |
| | | HSN (2) | | |
| | | PVC (2) | | |
| | | PVN (2) | | |
| | | PVP (2) | | |
| | | PVQ (2) | | |
| | | RID (1) | | |
| | | RIF (2) | | |
