## Supplementary figures and images for "Structural and developmental principles of neuropil assembly in *C. elegans*"

### Extended Data Figure 2

Extended Figure 2: Moyle et al.

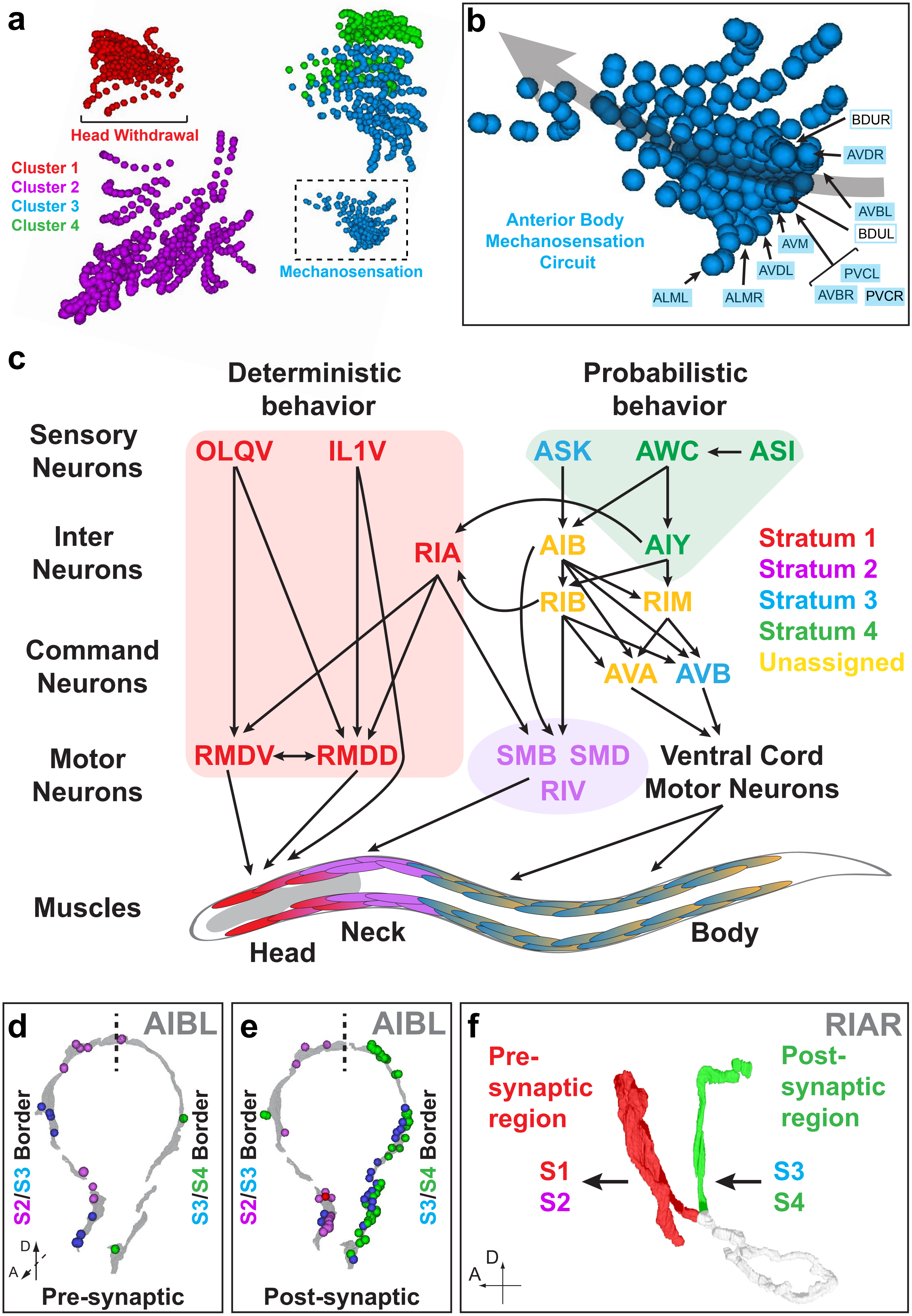

### Extended Data Figure 3

Extended Figure 3: Moyle et al.

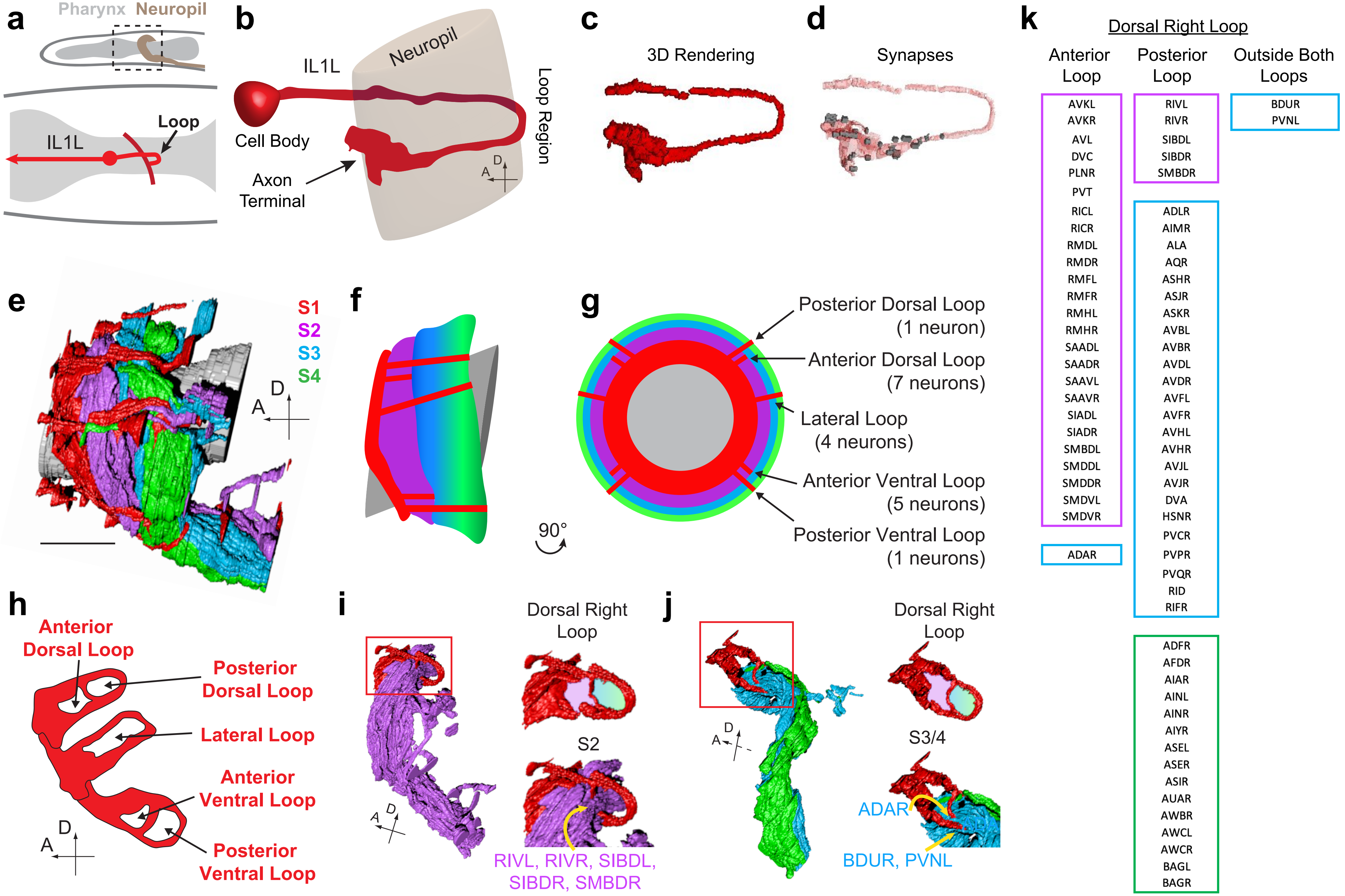

### Extended Data Figure 4

Extended Figure 4: Moyle et al.

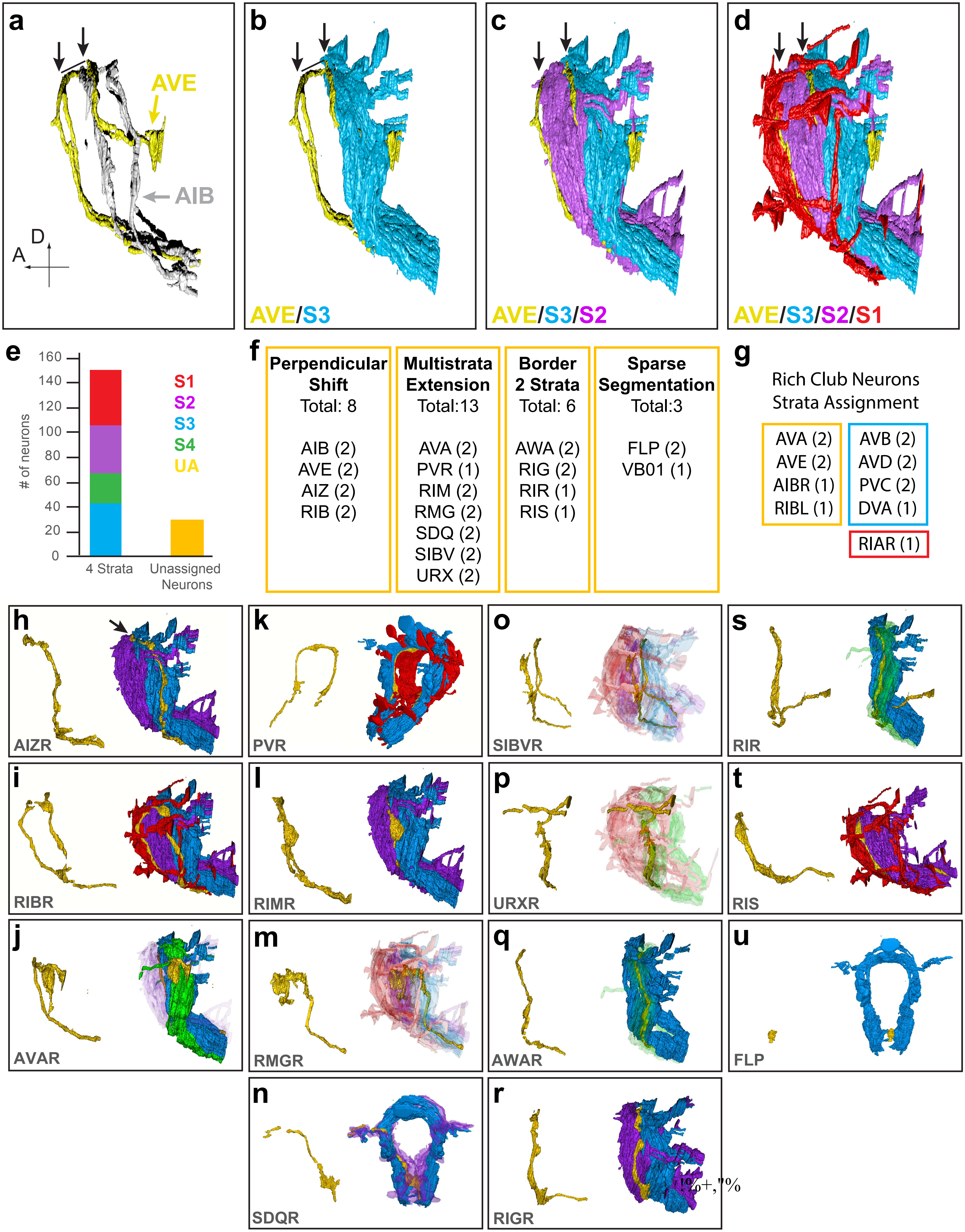

### Extended Data Figure 5

Extended Figure 5: Moyle et al.

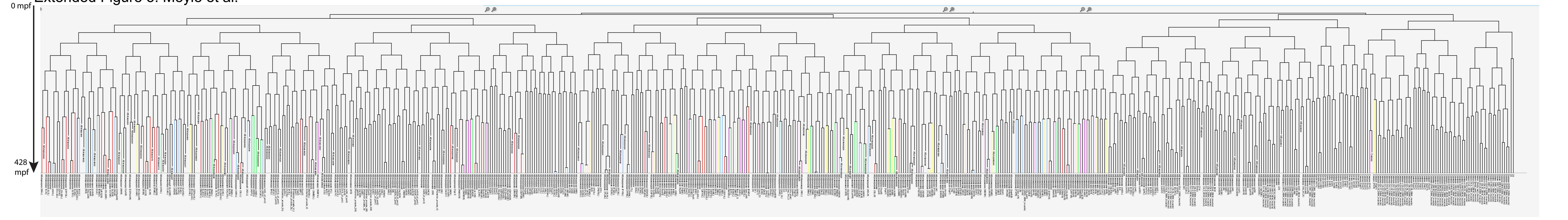

### Extended Data Figure 6

Extended Figure 6: Moyle et al.

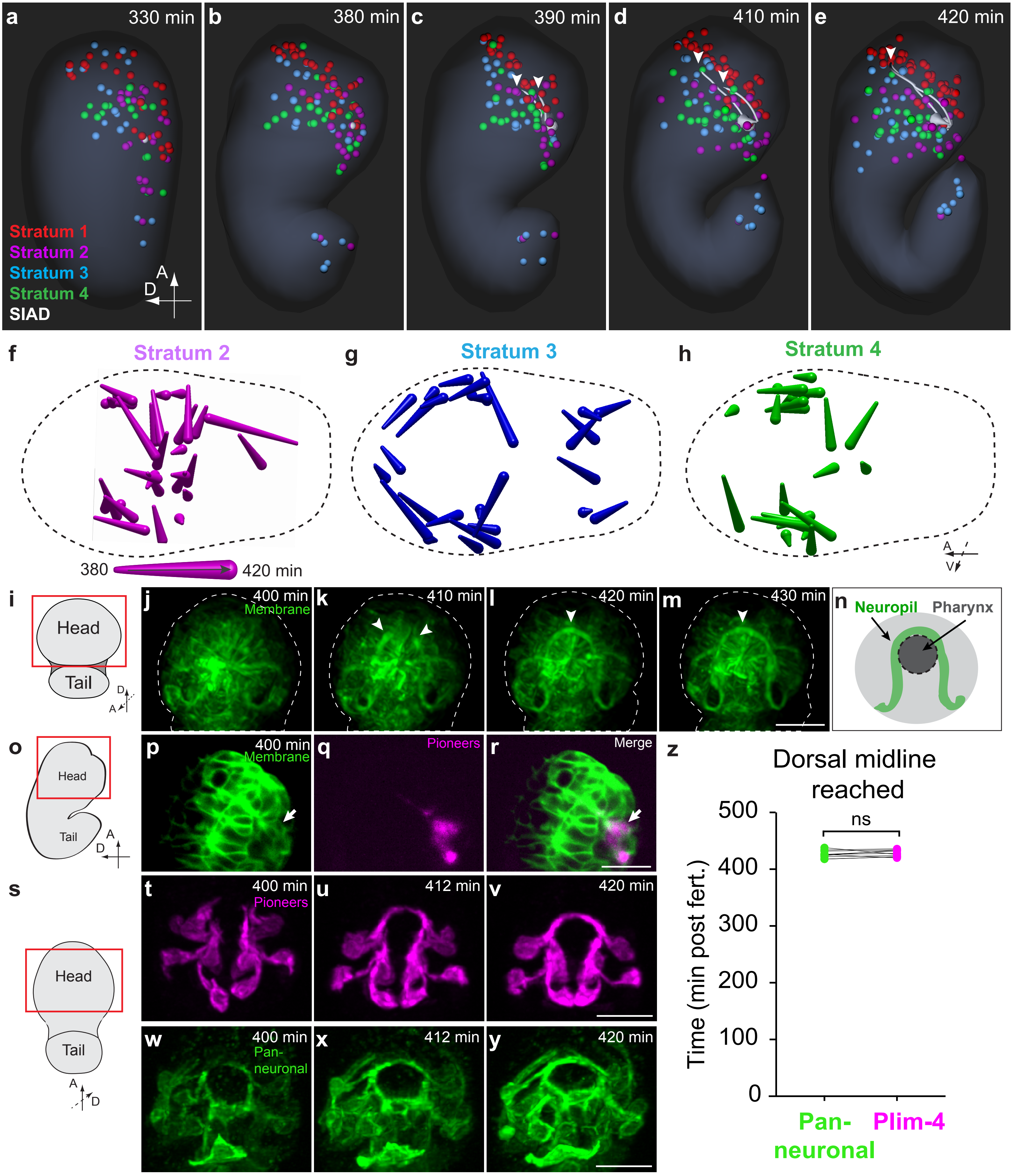

### Extended Data Figure 7

Extended Figure 7: Moyle et al.

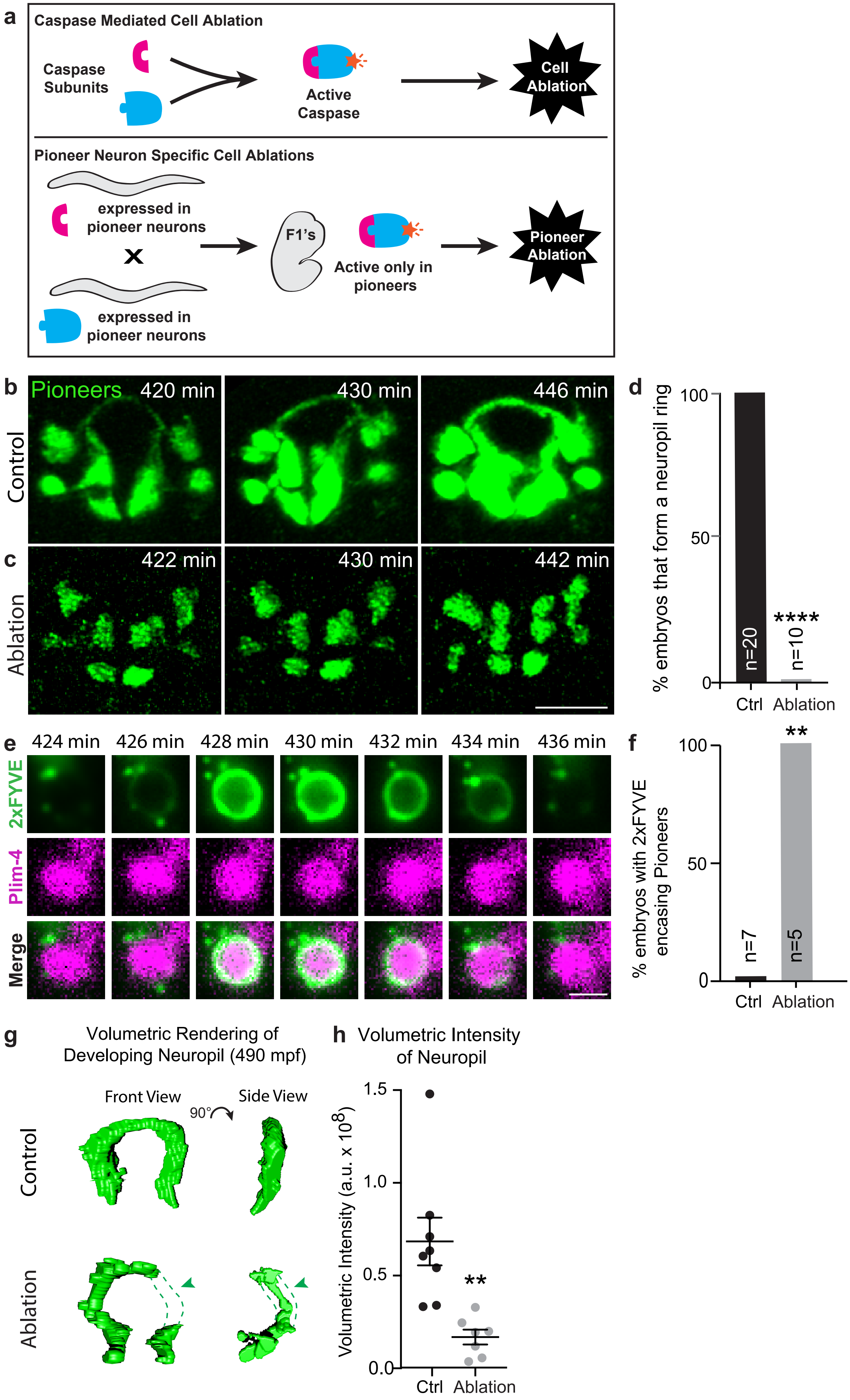

### Extended Data Figure 8

#### Extended Figure 8: Moyle et al.

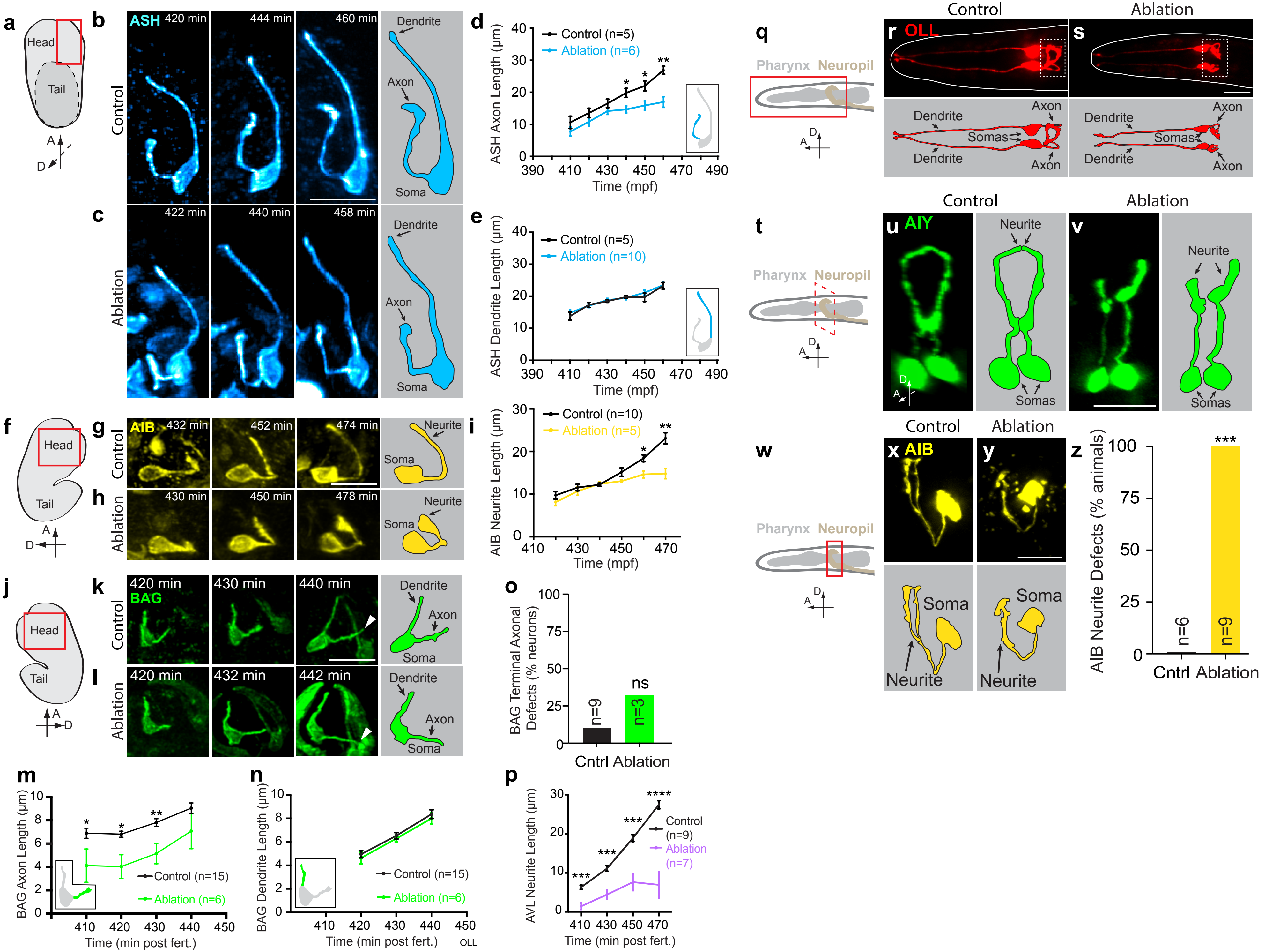

### Extended Data Figure 9

Extended Figure 9: Moyle et al.

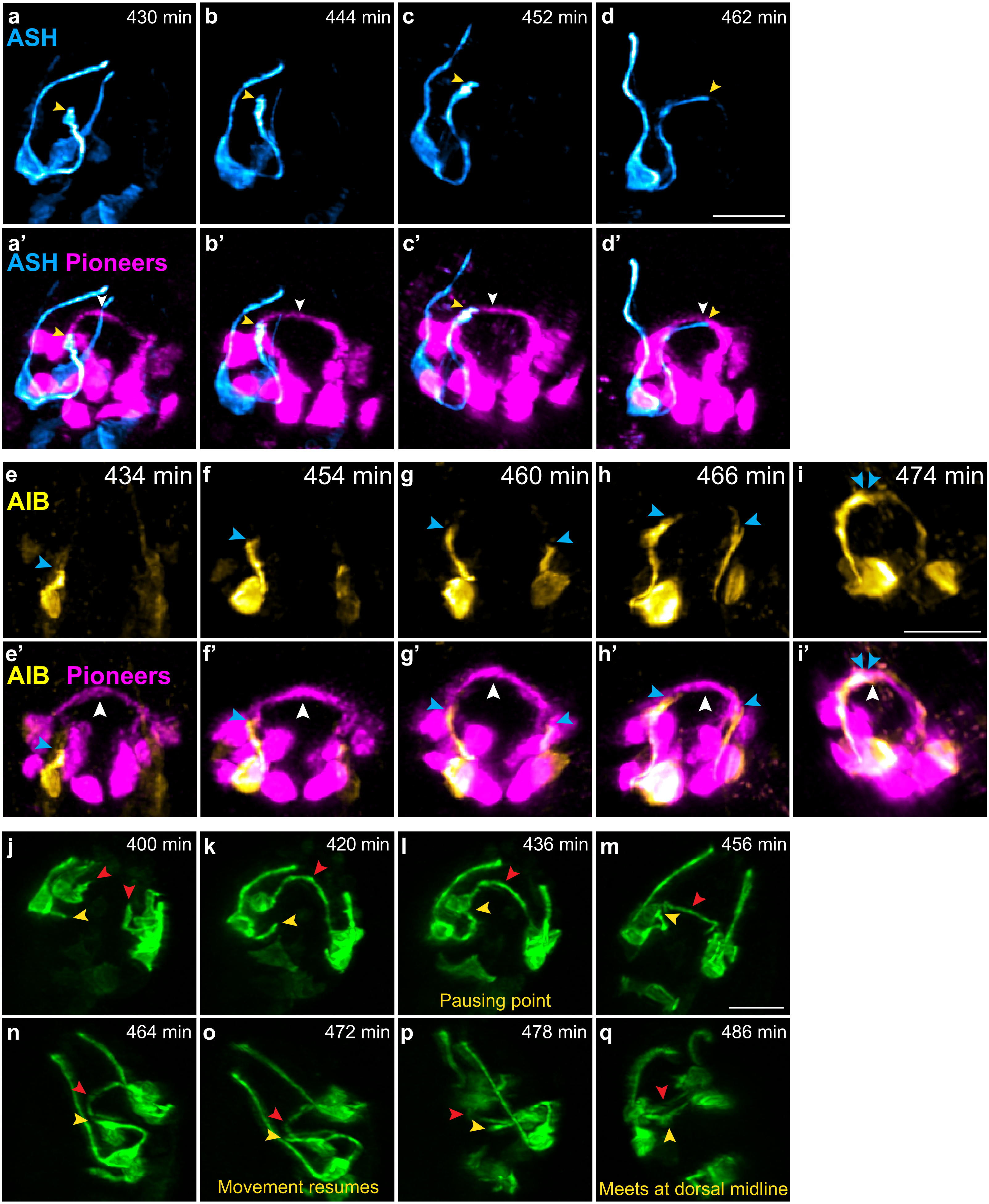
