## Supplemental Table 1 for "Structural and developmental principles of neuropil assembly in *C. elegans*"

Neurons colored according to stratum assignment:

| Stratum 2 | Stratum 3 | Stratum 4 |
|---|---|---|

### Dorsal Left

| Anterior Loop | Posterior Loop | Outside Both Loops |
|---|---|---|
| AVKL | RIVL | BDUL |
| AVKR | RIVR | PVNL |
| AVL | SIBDL | RIFL |
| DVC | SMBDL | |
| PLNL | ADFL | |
| PVT | AFDL | |
| RICL | AIAL | |
| RICR | AINL | |
| RMDL | AINR | |
| RMDR | AIYL | |
| RMFL | ASEL | |
| RMFR | ASER | |
| RMHL | ASIL | |
| RMHR | AUAL | |
| SAADL | AWBL | |
| SAADR | AWCL | |
| SAAVL | AWCR | |
| SAAVR | BAGL | |
| SIADL | BAGR | |
| SIADR | ADLL | |
| SIBDR | AIML | |
| SMBDR | ALA | |
| SMDDL | AQR | |
| SMDDR | ASHL | |
| SMDVL | ASJL | |
| SMDVR | ASKL | |
| ADAL | AVBL | |
| | AVBR | |
| | AVDL | |
| | AVDR | |
| | AVFL | |
| | AVFR | |
| | AVHL | |
| | AVHR | |
| | AVJL | |
| | AVJR | |
| | DVA | |
| | PVCL | |
| | PVCR | |
| | PVPL | |
| | PVQL | |

### Dorsal Right

| Anterior Loop | Posterior Loop | Outside Both Loops |
|---|---|---|
| AVKL | RIVL | BDUR |
| AVKR | RIVR | PVNL |
| AVL | SIBDL | |
| DVC | SIBDR | |
| PLNR | SMBDR | |
| PVT | ADFR | |
| RICL | AFDR | |
| RICR | AIAR | |
| RMDL | AINL | |
| RMDR | AINR | |
| RMFL | AIYR | |
| RMFR | ASEL | |
| RMHL | ASER | |
| RMHR | ASIR | |
| SAADL | AUAR | |
| SAADR | AWBR | |
| SAAVL | AWCL | |
| SAAVR | AWCR | |
| SIADL | BAGL | |
| SIADR | BAGR | |
| SMBDL | ADLR | |
| SMDDL | AIMR | |
| SMDDR | ALA | |
| SMDVL | AQR | |
| SMDVR | ASHR | |
| ADAR | ASJR | |
| | ASKR | |
| | AVBL | |
| | AVBR | |
| | AVDL | |
| | AVDR | |
| | AVFL | |
| | AVFR | |
| | AVHL | |
| | AVHR | |
| | AVJL | |
| | AVJR | |
| | DVA | |
| | HSNR | |
| | PVCR | |
| | PVPR | |
| | PVQR | |
| | RID | |
| | RIFR | |

### Lateral Left

| Inside Loop | Outside Loop |
|---|---|
| ALNL | ADEL |
| AVKL | ALML |
| AVKR | BDUL |
| AVL | PVNL |
| DVC | RIFL |
| PVT | |
| RICL | |
| RICR | |
| RIVL | |
| RIVR | |
| RMDL | |
| RMDR | |
| RMFL | |
| RMFR | |
| RMHL | |
| RMHR | |
| SAADL | |
| SAAVL | |
| SAAVR | |
| SIADL | |
| SIAVL | |
| SIBDL | |
| SMBDL | |
| SMBVL | |
| SMDDL | |
| SMDVL | |
| SMDVR | |
| ADFL | |
| AFDL | |
| AIAL | |
| AINL | |
| AIYL | |
| ASEL | |
| ASER | |
| ASIL | |
| AUAL | |
| AWBL | |
| AWCL | |
| AWCR | |
| BAGL | |
| BAGR | |
| ADAL | |
| ADLL | |
| AIML | |
| ALA | |
| AQR | |
| ASHL | |
| ASJL | |
| ASKL | |
| AVBL | |
| AVBR | |
| AVDR | |
| AVFL | |
| AVFR | |
| AVHR | |
| AVJR | |
| DVA | |
| PVCL | |
| PVCR | |
| PVPL | |
| PVQL | |

### Lateral Right

| Inside Loop | Outside Loop |
|---|---|
| ALNR | ADER |
| AVKL | ALMR |
| AVKR | BDUR |
| AVL | HSNR |
| DVC | PVNL |
| PVT | RIFR |
| RICL | |
| RICR | |
| RIVL | |
| RIVR | |
| RMDL | |
| RMDR | |
| RMFL | |
| RMFR | |
| RMHL | |
| RMHR | |
| SAADR | |
| SAAVL | |
| SAAVR | |
| SIADL | |
| SIADR | |
| SIAVR | |
| SIBDR | |
| SMBDR | |
| SMBVR | |
| SMDDR | |
| SMDVL | |
| SMDVR | |
| ADFR | |
| AFDR | |
| AIAR | |
| AINR | |
| AIYR | |
| ASEL | |
| ASER | |
| ASIR | |
| AUAR | |
| AWBR | |
| AWCL | |
| AWCR | |
| BAGL | |
| BAGR | |
| ADAR | |
| ADLR | |
| AIMR | |
| ALA | |
| AQR | |
| ASHR | |
| ASJR | |
| ASKR | |
| AVBL | |
| AVBR | |
| AVDL | |
| AVFL | |
| AVFR | |
| AVHL | |
| AVJL | |
| DVA | |
| PVCR | |
| PVPR | |
| PVQR | |
| RID | |

### Ventral Left

| Anterior Loop | Posterior Loop | Outside Both Loops |
|---|---|---|
| AVKL | SMBVL | BDUL |
| AVKR | ADFL | PVNL |
| AVL | AFDL | RIFL |
| DVC | AIAL | |
| PVT | AINL | |
| RICL | AIYL | |
| RICR | ASEL | |
| RIVR | ASGL | |
| RMDR | ASIL | |
| RMFL | AUAL | |
| RMFR | AWBL | |
| RMHL | AWCL | |
| SAADL | AWCR | |
| SAAVR | BAGR | |
| SIADL | ADLL | |
| SIAVL | AIML | |
| SIBDL | ALML | |
| SMBDL | AQR | |
| SMBVL | ASHL | |
| SMDDL | ASJL | |
| SMDVR | ASKL | |
| ADAL | AVBL | |
| | AVBR | |
| | AVDR | |
| | AVFL | |
| | AVFR | |
| | AVHR | |
| | AVJR | |
| | DVA | |
| | PVCL | |
| | PVPL | |
| | PVQL | |

### Ventral Right

| Anterior Loop | Posterior Loop | Outside Both Loops |
|---|---|---|
| AVKL | SMBVR | BDUR |
| AVKR | ADFR | HSNR |
| AVL | AFDR | PVNL |
| DVC | AIAR | RIFR |
| PVT | AINR | |
| RICL | AIYR | |
| RICR | ASER | |
| RIVL | ASGR | |
| RMDL | ASIR | |
| RMFR | AUAR | |
| RMHR | AWBR | |
| SAADR | AWCL | |
| SAAVL | AWCR | |
| SIADR | BAGL | |
| SIAVR | ADLR | |
| SIBDR | AIMR | |
| SMBDR | ALMR | |
| SMBVR | AQR | |
| SMDDR | ASHR | |
| SMDVL | ASJR | |
| ADAR | ASKR | |
| | AVBL | |
| | AVBR | |
| | AVDL | |
| | AVFL | |
| | AVFR | |
| | AVHL | |
| | AVJL | |
| | DVA | |
| | PVCR | |
| | PVPR | |
| | PVQR | |
| | RID | |
