## Supplemental Table 2 for "Structural and developmental principles of neuropil assembly in *C. elegans*"

| Strain Name: | Genotype: | Source: | Structures Labeled |
|---|---|---|---|
| N2 (ancestral) | | Caenorhabditis Genetics Center | |
| RW10226 | unc-119(ed3) III; itIs37 [pie-1p::mCherry::H2B::pie-1 3'UTR + unc-119(+)] IV; stIs10226 [his-72p::HIS-24::mCherry::let-858 3' UTR + unc-119(+)]. | Caenorhabditis Genetics Center | Nuclei |
| BV514 | ujIs113 [pie-1p::mCherry::H2B::pie-1 3'UTR + nhr-2p::his-24::mCherry::let-858 3'UTR + unc-119(+)]; II | Duncan et al., 2019 | Nuclei |
| DCR4111 | olaex2388 [DACR2257 at 10ng/uL + DACR218 at 30ng/uL]; unc-119(ed3) III; itIs37 [pie-1p::mCherry::H2B::pie-1 3'UTR + unc-119(+)] IV; stIs10226 [his-72p::HIS-24::mCherry::let-858 3' UTR + unc- | This Paper | All Membranes, Nuclei |
| DCR3969 | olaEx2302 [DACR2266 at 25ng/uL + DACR218 at 30ng/uL] | This Paper | 16 Pioneers |
| DCR3970 | olaEx2303 [DACR2266 at 25ng/uL + DACR218 at 30ng/uL] | This Paper | 16 Pioneers |
| DCR3971 | olaEx2304 [DACR2266 at 25ng/uL + DACR218 at 30ng/uL] | This Paper | 16 Pioneers |
| DCR6098 | olaIs93 [DACR2291 50 ng/uL + DACR2351 25 ng/uL + DACR218 30 ng/uL] X | This Paper | 16 Pioneers |
| DCR6100 | olaIs90 [DACR2291 50 ng/uL + DACR2351 25 ng/uL + DACR218 30 ng/uL] X | This Paper | 16 Pioneers |
| DCR6102 | olaIs85 [DACR2292 at 50 ng/uL + DACR20 at 25 ng/uL] IV | This Paper | |
| DCR6106 | olaIs85 [DACR2292 at 50 ng/uL + DACR20 at 25 ng/uL] IV | This Paper | 16 Pioneers |
| DCR6107 | olaIs85 [DACR2292 at 50 ng/uL + DACR20 at 25 ng/uL] IV | This Paper | 16 Pioneers |
| DCR6269 | olaIs85 [DACR2292 at 50 ng/uL + DACR20 at 25 ng/uL] IV | This Paper | Pan-neuronal |
| DCR6186 | olaIs85 [DACR2292 at 50 ng/uL + DACR20 at 25 ng/uL] IV | This Paper | Pan-neuronal |
| OH812 | olaIs85 [DACR2292 at 50 ng/uL + DACR20 at 25 ng/uL] IV | Caenorhabditis Genetics Center | AVL |
| DCR6975 | olaIs85 [DACR2292 at 50 ng/uL + DACR20 at 25 ng/uL] IV | This Paper | AVL |
| DCR7856 | olaIs85 [DACR2292 at 50 ng/uL + DACR20 at 25 ng/uL] IV | This Paper | OLL in L1s; Pioneers in L1s |
| DCR7871 | olaIs85 [DACR2292 at 50 ng/uL + DACR20 at 25 ng/uL] IV | This Paper | OLL in L1s |
| DCR6853 | olaIs85 [DACR2292 at 50 ng/uL + DACR20 at 25 ng/uL] IV | This Paper | AIB, ASH; Nuclei |
| DCR6631 | olaIs85 [DACR2292 at 50 ng/uL + DACR20 at 25 ng/uL] IV | This Paper | AIB, ASH |
| DCR6633 | olaIs85 [DACR2292 at 50 ng/uL + DACR20 at 25 ng/uL] IV | This Paper | AIB, ASH |
| DCR6182 | olaIs85 [DACR2292 at 50 ng/uL + DACR20 at 25 ng/uL] IV | This Paper | AIB, ASH, Pioneers |
| DCR7844 | olaIs85 [DACR2292 at 50 ng/uL + DACR20 at 25 ng/uL] IV | This Paper | AIY in L1s; Pioneers in L1 |
| DCR7870 | olaIs85 [DACR2292 at 50 ng/uL + DACR20 at 25 ng/uL] IV | This Paper | AIY in L1s |
| DCR6358 | olaIs85 [DACR2292 at 50 ng/uL + DACR20 at 25 ng/uL] IV | This Paper | BAG; Nuclei |
| DCR6652 | olaIs85 [DACR2292 at 50 ng/uL + DACR20 at 25 ng/uL] IV | This Paper | BAG |
| DCR6653 | olaIs85 [DACR2292 at 50 ng/uL + DACR20 at 25 ng/uL] IV | This Paper | BAG |
| DCR6744 | olaIs85 [DACR2292 at 50 ng/uL + DACR20 at 25 ng/uL] IV | This Paper | BAG |
| DCR7692 | olaIs85 [DACR2292 at 50 ng/uL + DACR20 at 25 ng/uL] IV | Duncan et al., 2019 | RMDD |
| DCR3985 | olaIs85 [DACR2292 at 50 ng/uL + DACR20 at 25 ng/uL] IV | This Paper | All Membranes |
| DCR3986 | olaIs85 [DACR2292 at 50 ng/uL + DACR20 at 25 ng/uL] IV | This Paper | All Membranes |
| DCR6142 | olaIs85 [DACR2292 at 50 ng/uL + DACR20 at 25 ng/uL] IV | This Paper | All Membranes; 16 Pioneers |
| DCR6143 | olaIs85 [DACR2292 at 50 ng/uL + DACR20 at 25 ng/uL] IV | This Paper | All Membranes; 16 Pioneers |
| DCR4485 | olaIs85 [DACR2292 at 50 ng/uL + DACR20 at 25 ng/uL] IV | This Paper | Pan-neuronal |
| DCR6139 | olaIs85 [DACR2292 at 50 ng/uL + DACR20 at 25 ng/uL] IV | This Paper | Pan-neuronal; 16 Pioneers |
| DCR6140 | olaIs85 [DACR2292 at 50 ng/uL + DACR20 at 25 ng/uL] IV | This Paper | Pan-neuronal; 16 Pioneers |
| DCR6861 | olaIs85 [DACR2292 at 50 ng/uL + DACR20 at 25 ng/uL] IV | This Paper | Cell Death Marker; 16 Pioneers |
| DCR6864 | olaIs85 [DACR2292 at 50 ng/uL + DACR20 at 25 ng/uL] IV | This Paper | Cell Death Marker; 16 Pioneers |
| DCR6862 | olaIs85 [DACR2292 at 50 ng/uL + DACR20 at 25 ng/uL] IV | This Paper | Cell Death Marker |
| DCR7862 | olaIs85 [DACR2292 at 50 ng/uL + DACR20 at 25 ng/uL] IV | This Paper | AIB in L1s; Pioneers in L1s |
| DCR7872 | olaIs85 [DACR2292 at 50 ng/uL + DACR20 at 25 ng/uL] IV | This Paper | AIB in L1s |
| DCR6298 | olaIs85 [DACR2292 at 50 ng/uL + DACR20 at 25 ng/uL] IV | This Paper | SAAV, AFD, ADF, AWC, AWB |
