## Supplemental Table 3 for "Structural and developmental principles of neuropil assembly in *C. elegans*"

| Plasmid Name | Promoter (gene) | Size (bp) | Oligonucleotide 1 (5 → 3) | Oligonucleotide 2 (5 → 3) | Full Construct |
|---|---|---|---|---|---|
| DACR2257 | nhr-2 | 2418 | CTCGTCGAGAAGGCATACAGTAG | CTGAAAACTAGAAAAGAAATAGAGATAGAAAAATAATAAG | Pnhr-2::PHD::GFP::unc54UTR |
| DACR2301 | rab-3 | 4383 | GCGAGTTTTGACTGGCTTTC | CTGAAAATAGGGCTACTGTAGATTTATTTTAAAAG | Prab-3(includes Exon1)::SL2::PHD::GFP::unc-54UTR |
| DACR2292 | lim-4 | 4605 | CCCATGCAGTTCAAATACTGTC | CGGAAATAACATCCTTTACAGTAAAACG | Plim-4(includes Exons 1-3)::SL2::p17-Caspase3::unc-54UTR |
| DACR2291 | lim-4 | 4605 | CCCATGCAGTTCAAATACTGTC | CGGAAATAACATCCTTTACAGTAAAACG | Plim-4(includes Exons 1-3)::SL2::p12-Caspase3::unc-54UTR |
| DACR2280 | ceh-24 | 2900 | GAACACCATCGCTCTCTCATC | ACTTCCAAGGCAGAGAGCTG | Pceh-24::PHD::GFP::unc-54UTR |
| DACR2266 | lim-4 | 4605 | CCCATGCAGTTCAAATACTGTC | CGGAAATAACATCCTTTACAGTAAAACG | Plim-4(includes Exons 1-3)::PHD::GFP::unc-54UTR |
| DACR3254 | ceh-24 | 2900 | GAACACCATCGCTCTCTCATC | ACTTCCAAGGCAGAGAGCTG | Pceh-24::mCherry::unc-54UTR |
| DACR2609 | unc-42 | 3176 | GTCCTGTCGATGCCATTTTTGTG | TGTGTGAGTGAAAGCGGAGA | Punc-42::ZF1::PHD::GFP::unc-54UTR |
| DACR2607 | lim-4 | 4605 | CCCATGCAGTTCAAATACTGTC | CGGAAATAACATCCTTTACAGTAAAACG | Plim-4(includes Exons 1-3)::SL2::ZIF-1::unc-54UTR |
| DACR2436 | elt-7 | 2414 | ATGCAACACAGAGAACACTTCC | TTTTTCCAGTCGACTAGAGCAGAC | Pelt-7::mCherry::NLS::unc-54UTR |
| DACR2410 | elt-7 | 2414 | ATGCAACACAGAGAACACTTCC | TTTTTCCAGTCGACTAGAGCAGAC | Pelt-7::GFP::NLS::unc-54UTR |
| DACR2371 | unc-42 | 3176 | GTCCTGTCGATGCCATTTTTGTG | TGTGTGAGTGAAAGCGGAGAAATG | Punc-42::PHD::GFP::unc-54UTR |
| DACR2351 | lim-4 | 4605 | CCCATGCAGTTCAAATACTGTC | CGGAAATAACATCCTTTACAGTAAAACG | Plim-4(includes Exons 1-3)::mCherry::unc-54UTR |
| DACR2605 | egl-13 | 6513 | CCGATTTGGCCAATGGTACTTG | CAGGACCAAGAATGGACAACTCAC | Pegl-13::egl-13(168bp)::SL2::PHD::unc-54UTR |
| DACR2550 | ceh-48 | 2617 | CAGAGGCAGAAGCTCGAAAGCATG | CTCCAAGGGTCCTCCTGAAAATG | Pceh-48(2616bp)::artificial_intron::SL2::PHD::GFP::unc-54UTR |
| DACR2603 | sdz-31 | 1373 | GCATCGTTGTGCAGTTGGAATTC | TAAACCTCGTGGCCACGCC | Psdz-31::GFP::NLS::unc-54UTR |
| DACR2541 | ceh-37 | 3259 | TTGATACTTACGCGTGTCCGACAG | CTGCATTGTATGGTGCTCCACTAAAC | Pceh-37::ceh-37(103bp)::SL2::PHD::GFP::unc-54UTR |
| DACR2505/pNL1 | ced-1 | 5076 | Lu et al., 2009 | Lu et al., 2009 | Pced-1::2xFYVE::GFP(S65C,Q80R)::unc-54UTR |
| DACR2245 | inx-1 | 1000 | ATTATTTTCTGTGCTTTTCACAAATACAC | TCCGGCGGACAAGAAC | Pinx-1::eGFP::Rab-3::unc-54UTR |
| DACR1412 | inx-1 | 451 | TATAGTTCTTCATCTTCTTTTTTTTAATATCCTC | TCCGGCGGACAAGAAC | Pinx-1::mCherry::unc-54UTR |

PHD: Pleckstrin homology domain used for membrane targeting
SL2: Trans-spliced RNA leader sequence 2 in *C. elegans*
NLS: Nuclear localization signal
